# Using population dynamics to count bacteriophages and their lysogens

**DOI:** 10.1101/2023.10.06.561271

**Authors:** Yuncong Geng, Thu Vu Phuc Nguyen, Ehsan Homaee, Ido Golding

**Author notes:** Y.G. and T.V.P.N. contributed equally to this work.

## Abstract

Traditional assays for counting bacteriophages and their lysogens are labor-intensive and highly perturbative to the host cells. Here, we present a high-throughput infection method where all steps—cell growth, viral encounters, and post-infection recovery—take place in a microplate reader, and the growth dynamics of the infected culture are measured continuously using the optical density (OD). We find that the post-infection dynamics are reproducible and interpretable. In particular, the OD at which the culture lyses scales linearly with the logarithm of the initial phage concentration, providing a way of measuring phage numbers in unknown samples over nine decades and down to single-phage sensitivity. Interpreting the measured dynamics using a mathematical model for the coupled kinetics of phages and bacteria further allows us to infer the rates of viral encounters and cell lysis. Adding a single step of antibiotic selection provides the ability to measure the rate of host lysogenization. To demonstrate the application of our assay, we characterized the effect of bacterial growth rate on the propensity of lambda phage to lysogenize *E. coli*. When infected by a single phage, the probability of lysogenization is found to decrease approximately exponentially with the host growth rate. In growing, but not in stationary, cells, the propensity to lysogenize increases ~50-fold when multiple phages co-infect the cell. These findings illuminate how host physiology feeds into the lysis/lysogeny decision circuit, and demonstrate the utility of high-throughput infection to interrogating phage-host interactions.

---

An essential element in laboratory studies of bacteriophages is the counting of phages and of cells undergoing lysogeny. The protocols for performing these tasks typically consist of the following steps: (i) pre-infection cell growth, (ii) incubation for phage adsorption, (iii) triggering phage genome injection, (iv) post-infection cell recovery, and (v) measurement of the infection outcome^1–3^. The procedure involves centrifugation, incubation without aeration, and temperature changes, thus strongly perturbing the pre-infection cellular state. Consequently, the impact of host physiology on infection outcome—often of significant interest^4–6^—is hard to evaluate. Furthermore, measuring this outcome typically relies on plating and requires multiple dilutions to obtain countable numbers of plaques or colonies. These low-throughput steps hinder scaling up the experiments, in turn limiting the ability to perform systematic sampling of parameters of interest.

To overcome these deficiencies, we devised a high-throughput assay (**Figure 1A**) where phage infection takes place under uninterrupted cell growth in a microplate reader, and the infection outcome is monitored using the culture’s growth dynamics, read continuously from the optical density (OD). Multiple samples under different infection conditions, e.g., multiplicity of infection (MOI) or growth media, can be assayed simultaneously. The post-infection growth dynamics can be used to estimate the number of phages in an unknown sample. Interpreted using a model for the coupled kinetics of bacterial and viral populations, the measured dynamics further allow inferring the phage encounter rate, latent period, and burst size. Finally, adding a single step of antibiotic selection provides the ability to measure the probability of host lysogenization.

**Figure 1:**
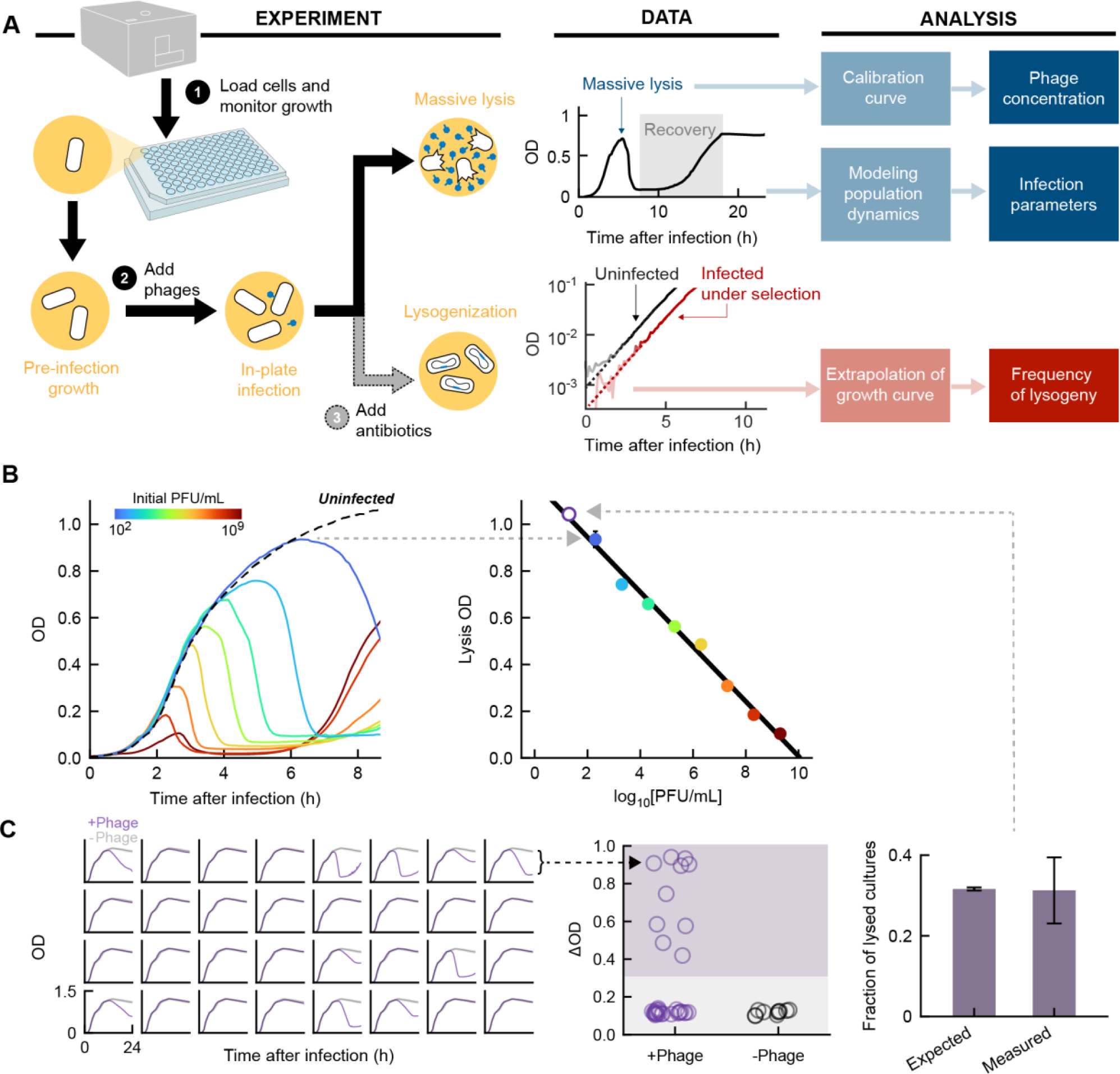
A high-throughput method for counting phages and lysogenic cells in a microplate reader. **(A)** Experimental and analytical pipeline. Left, bacterial cultures are grown and infected in a microwell plate reader, where the optical density (OD) is continuously measured. To count phages, no additional experimental manipulation is needed. To measure the frequency of lysogeny, a step of antibiotic selection is performed. Middle, growth curves of *E. coli* cultures, infected by (top) obligately lytic phage (λ_ts_ at 37°C), and (bottom) temperate phage (λ_ts_ at 30°C) under antibiotic selection. Right: Phage concentrations, infection parameters, and frequency of lysogeny are inferred from different features of the growth dynamics. **(B)** OD-based phage counting. Left, solid lines, growth curves of *E. coli* MG1655 cultures at 37°C in LBM infected by λ_ts_ at different concentrations (different colors). Dashed line, growth curve of an uninfected culture. Right, the OD at which the culture lyses scales linearly with the logarithm of the initial phage concentrations over 9 orders of magnitude. Colored markers, data; error bars, standard errors of the mean (SEM) from four culture replicates. Empty marker, phages at very low numbers are counted using the modified assay shown in Panel C. Black line, linear fit. **(C)** Single-phage sensitivity of the counting assay. Left, purple lines, growth curves of 32 cell cultures, each infected by ≈ 0.4 PFU, at 37°C in LBM supplemented with 0.2% maltose. In each subplot, the growth curve of an uninfected sample (averaged over 10 replicates) is shown in gray. Middle, the difference in OD between the first local maximum and the subsequent minimum for each culture (denoted ΔOD). Cultures are considered lysed if ΔOD > 0.3 (purple shading). Right, the expected fraction of wells with non-zero phage numbers, estimated from plating, and the measured fraction of lysed cultures. Error bars, SEM.

We developed our protocol using bacteriophage lambda, owing to the system’s incomparable knowledge base^7^ and our lab’s experience with it^8–10^. Pre-infection *E. coli* cultures (MG1655) were grown in LBM (LB supplemented with 10 mM MgSO_4_) in microplate wells at constant temperature (37°C) and aeration, and samples of phage (λ *cI*857 *bor*::*kan*^R^, hereafter λ_ts_, obligately lytic at 37°C, ref. ^11^) were directly added to the cultures during exponential phase (See ***SI Methods***). The absorbance at 595 nm (optical density, OD_595_) of each culture, which can be converted to bacterial concentration^12^, was recorded by the microplate reader throughout the experiment. A typical OD curve is shown in **Figure 1A**. After an initial increase due to cell growth, a drop in bacterial density is observed, reflecting the well-documented phenomenon of massive lysis^13,14^. Following a pause, the OD rises again and eventually reaches saturation. The measured OD curves are highly reproducible across biological repeats (**Figure S1**). Curves with the same qualitative characteristics were obtained using several lambda and *E. coli* strains, various growth media, as well as phages T4, T5, and P1 (**Figure S2**; see **Table S1** for list of bacterial and phage strains used in this study).

When using the protocol above to infect bacteria at a given density by varying concentrations of phages, we observed that different initial conditions resulted in clearly distinguishable OD curves (**Figure 1B**). In particular, the OD at which massive lysis begins (denoted hereafter “lysis OD”) scales linearly with the logarithm of the initial phage concentration (**Figure 1B**). This linear scaling, which holds over 9 orders of magnitude, provides a simple means of counting phages: A calibration curve is first obtained using serial dilution of a phage sample with known concentration, and then used to convert the lysis OD of an unknown sample to its phage concentration (**Figure S1**); no dilution or plating is needed. Moreover, the assay is sensitive to the presence of even a single phage: When the average number of infecting phages per well was less than one, the fraction of lysed cultures matched the expected fraction of wells with non-zero phage numbers (**Figure 1C**; see ***SI Methods***). Thus, our phage counting protocol involves no cost in sensitivity as compared to traditional plaque plating. A monotonic relation between lysis OD and initial phage concentration was also found in other lambda strains (both obligately lytic and temperate, see below), other growth media, and in phages T4, T5, and P1 (**Figure S3**), suggesting that the method for phage counting is broadly applicable.

Motivated by the interpretive power of the lysis OD with regards to the initial phage numbers, we reasoned that additional infection parameters may be encoded by the entirety of the measured curve. To infer these parameters, we performed infection at six different MOIs. In addition to continuously following the bacterial OD, we also measured the phage kinetics in the same cultures by extracting samples at 10 time points and quantifying using the OD-based method described above (See ***SI Methods***). To interpret the data, we formulated a mathematical model describing the coupled dynamics of four species: nutrients (*N*), uninfected cells (*U*), infected cells (*I*), and phages (*P*), through three biological processes: cell growth, phage/cell encounters, and cell lysis^15^ (**Figure 2A**; see ***SI Methods***). We then proceeded to identify the associated kinetic schemes and parameters as follows.

i. Cell growth: We assumed that the instantaneous growth rate *g* depends on available nutrients via the Monod equation^16^: *g*(*N*) = *vN*/(*N* + *K*), where the single species *N* represents the pool of growth-limiting substrates in the medium. When multiple substrates are present, they are consumed sequentially, resulting in several growth phases, each characterized by specific values of the maximal growth rate *v* and Monod constant *K* (refs. ^17,18^). We used the growth curves of uninfected cells to infer *v* and *K* at each phase. As expected, growth in single-carbon media (M9 minimal broth supplemented with 0.4% glucose or 0.4% maltose) was describable using a single phase of nutrient consumption^19^ (**Figure S4** and **Table S2**), whereas the growth curves in complex media indicated successive consumption phases: two phases for tryptone broth supplemented with 10 mM MgSO_4_ (TBM, **Figure S4** and **Table S3**), and three phases for LBM (**Figure 2B** and **Table S4**).
ii. Cell lysis: Following refs. ^20,21^, we assumed that an infected cell goes through *M* intermediate states (*I*_1_, *I*_2_, …, *I*_*M*_) before lysis, with equal transition rates (= *M*/*τ*) from one state to the next. Consequently, the time between infection and cell lysis (the latent period) follows an Erlang distribution with mean *τ* and shape parameter *M*. We used the post-lysis dynamics of infected cultures to infer *M* and *τ* (**Figure 2C**). We found that the average latent period *τ* is well approximated by a linear function of the doubling time preceding massive lysis (**Figure 2C** and **Table S5**), a trend consistent with previous reports for phage T4 (ref. ^22^).
iii. To estimate the remaining parameters (the rate constant for successful phage/cell encounter *r* and the lytic burst size *B*), we fitted the model, simultaneously, to the cell densities and phage concentrations measured from all six infected cultures. Fitting was performed by minimizing the mean absolute error (MAE) using simulated annealing^23^.

**Figure 2.**
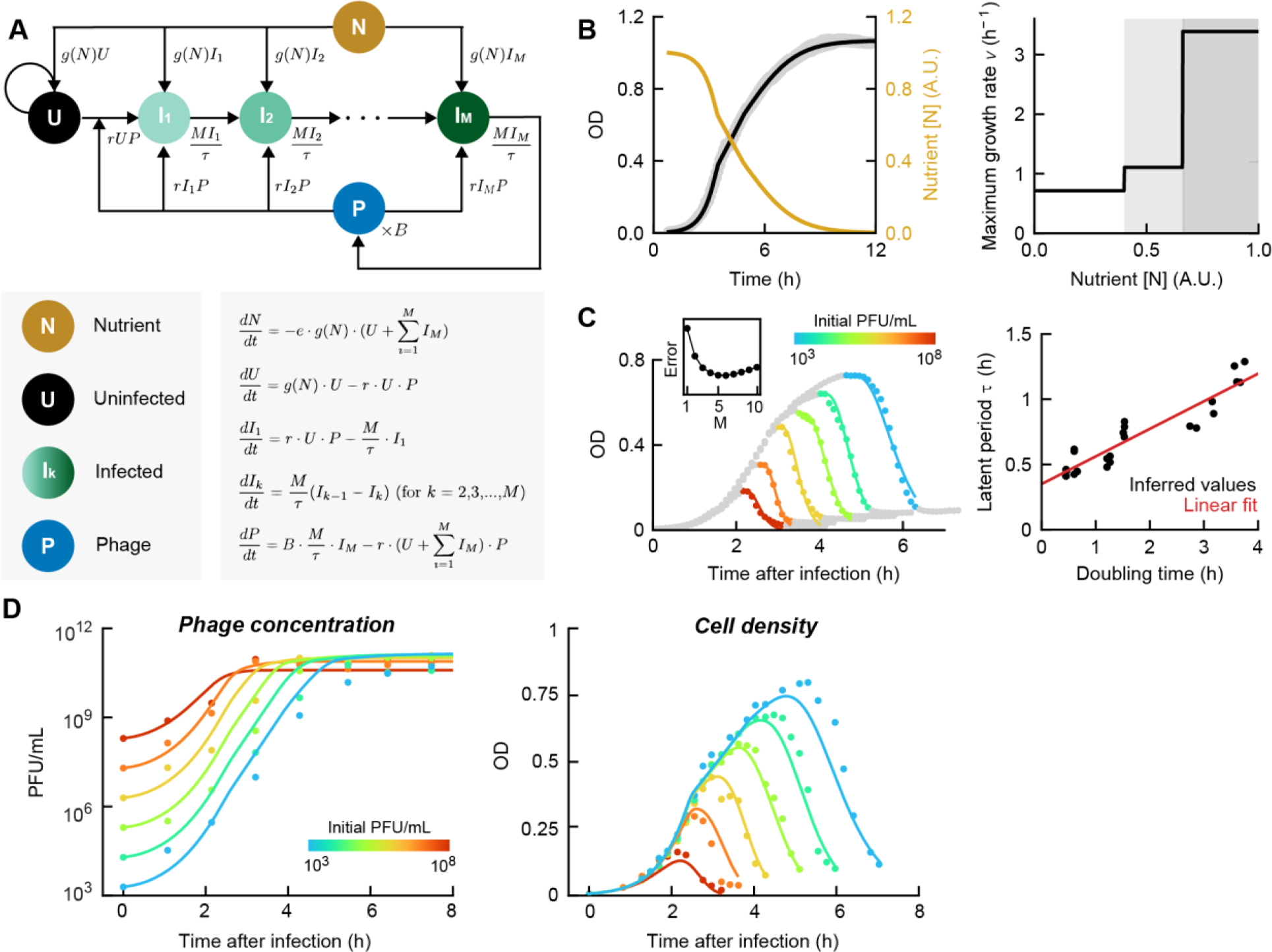
A mathematical model captures the growth dynamics of phages and bacteria and allows inference of infection parameters. **(A)** Model schematics and equations. Circles, species tracked by the model. Arrows, transitions between species. The transition rates are indicated next to the corresponding arrows. **(B)** Parameterization of bacterial growth. Left, a model describing nutrient-dependent growth (black) captures the OD curves of uninfected cultures (gray); gold, the inferred time-dependent nutrient abundance. Right, the maximum growth rate *v* at different stages of nutrient consumption (white and gray shading). **(C)** Parameterization of the latent period. Left, the latent period *τ* and shape parameter *M* are estimated from the postlysis growth dynamics. Markers, OD of cultures infected at different phage concentrations; markers are colored if used for fitting, and gray otherwise. Colored lines, model fits. Inset, the mean absolute error (MAE) as a function of the shape parameter *M*. Right, the mean latent period *τ*, inferred from the fits in the left panel, as a function of the doubling time preceding massive lysis. Black markers, fitted values. Red line, linear fit. **(D)** Model fitting for the measured phage concentrations (left) and cell densities (right) over time. Colored markers, data from infection at different initial phage concentrations. Colored lines, model fits.

The fitted model agreed well with the experimental data (**Figure 2D** and **Table S6**). The inferred parameter values for the burst size (*B =* 207 ± 4) and latent period (*τ* = 25.5 ± 1.6 min at early exponential growth) were consistent with reported values^11,24,25^ and with our experimental measurements using standard phage protocols (**Figure S5**; see ***SI Methods***), thus lending further credence to the model. We were also able to successfully apply a similar fitting procedure to infection data in other growth media and other phages (**Figure S6** and **Table S6**).

As noted above, the post-infection dynamics continue beyond the lytic collapse. One noticeable feature is the subsequent recovery of culture growth (**Figure 1A**). This recovery is observed for all phages examined, including λ_ts_ (obligately lytic at 37°C) and the virulent phages T4 and T5 (**Figure S2**), thus does not reflect the growth of lysogenic cells (discussed below). Rather, growth recovery likely reflects the emergence of a resistant population^26,27^. Consistent with this interpretation, adding to our model above a first-order transition from uninfected (sensitive) to resistant cells^27^ captured the observed growth recovery (**Figure S7**). The inferred rate of switching into resistance, (6.6 ± 0.8) × 10^−7^ per min, was comparable with a previously reported value^27^.

But whereas growth recovery was observed for both temperate and virulent phages, the degree of population collapse preceding recovery was markedly different in the two cases: Cultures infected by wild-type lambda phage (λ *cI*_wt_ *bor*::*kan*^*R*^, hereafter λ_wt_) showed a smaller drop in OD compared to those infected by an obligately lytic strain (λ_ts_ at 37°C, **Figure 3A**). We reasoned that the higher survival in cultures infected by wild-type phage reflects the presence of lysogenic cells, which then resume growth and are immune to further infections^28^. This interpretation was confirmed by the antibiotic resistance (*kan*^*R*^, conferred by the prophages) of the surviving cells following infection by λ_wt_ (**Figure S8**). Incorporating into our model the formation of lysogenic cells, via a constant lysogenization frequency per infection, allowed the model to capture the difference in post-lysis dynamics between virulent and temperate strains, but required fitting the lysogenization frequency independently for each experimental condition (**Figure S9**). This is unsurprising, considering that the single frequency coarse grains over the entire history of the infected culture, during which the infection conditions—e.g., growth rate and MOI— constantly change (**Figure S10**). Since these parameters are expected to influence the propensity to lysogenize^8,29,30^, it would be more informative to measure the occurrence of lysogeny after a single infection cycle, during which the infection parameters are well-defined.

**Figure 3:**
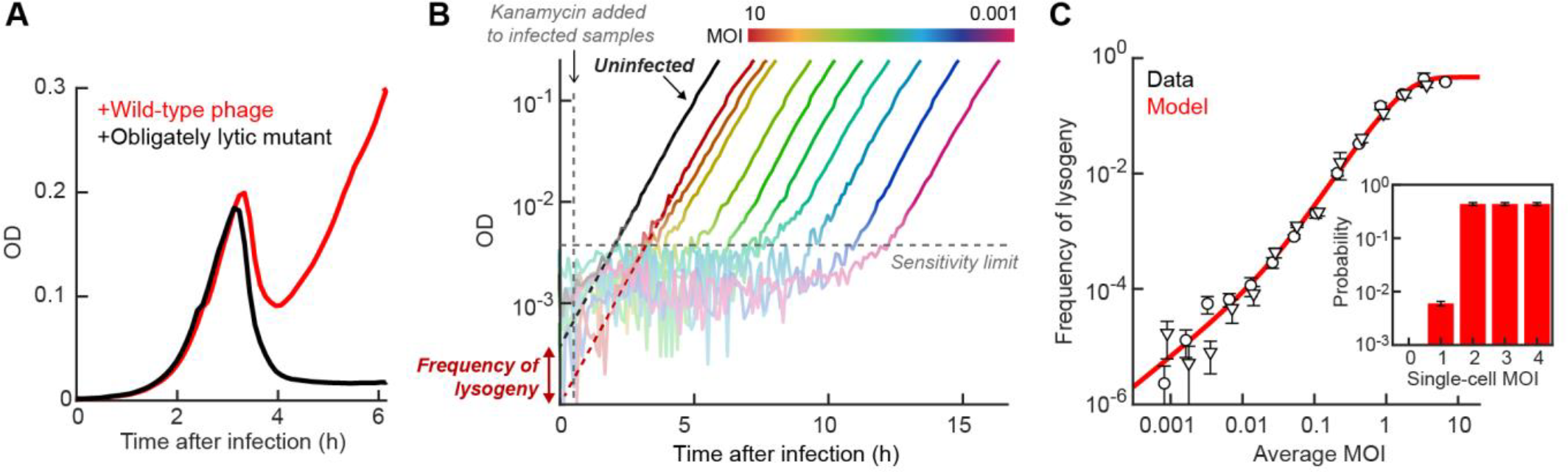
Imposing antibiotic selection after a single infection cycle allows counting of lysogens. **(A)** Difference in the population collapse between cultures infected by a temperate phage (λ_wt_, red) and obligately lytic mutant (λ_ts_, black). Cultures of *E. coli* MG1655, grown at 37°C in LBM, were both infected at MOI ≈ 10. **(B)** Using the growth dynamics under antibiotic selection to infer the fraction of cells undergoing lysogeny. Black line, OD of an uninfected culture without selection. Lines in other colors, OD of cultures infected by λ_ts_ at 30°C with different multiplicities of infection (MOIs), under kanamycin selection. All cultures were grown in LBM. Dashed lines, extrapolation of the OD back to *t* = 0, to infer the initial cell density. **(C)** The frequency of lysogeny as a function of MOI (adjusted for the fraction of phage-infected cells). Circles and triangles, data obtained in two independent runs of the experiment; error bars, SEM. Red line, model fit. Inset, the inferred probability of lysogenization as a function of the single-cell MOI.

To achieve this goal, we again utilized the lambda strain carrying an antibiotic resistance cassette (λ_ts_, *cI*857 *bor*::*kan*^*R*^). Following 15 minutes of microplate infection at 30°C (where λ_ts_ exhibits wild-type phenotype^11^), the culture was diluted 250-fold into fresh medium, reducing further infection^3,31^. After an additional 45 minutes of growth, kanamycin was added, allowing only lysogenic cells (carrying the resistance -encoding prophage) to grow^8,10,30^ (**Figure S11**; see ***SI Methods***). The growth curves under selective media were then extrapolated back to the time of dilution to infer the initial abundance of lysogens (**Figure 3B**). To validate our experimental approach, we used it to measure the frequency of lysogeny of MG1655 by λ_ts_ as a function of MOI (adjusted for the fraction of cells infected within 15 minutes; see ***SI Methods***). The measured curve (**Figure 3C**) recapitulates data obtained using a standard lysogenization protocol^10^. The data can be further used to infer the corresponding single-cell MOI response, by utilizing the Poissonian statistics of phage-cell encounters^29^. While similar inference was performed previously^8,10^, the higher throughput of the new protocol facilitates a finer sampling of MOI values, in turn constraining the single-cell relation better. Specifically, while earlier analysis indicated that infection by a minimum of two phages is required for lysogeny^10,29^, we were able to identify a non-zero probability of lysogenization during single-phage infection (**Figure 3C**, inset). This finding may help reconcile the bulk data with the gradual MOI response observed in single-cell experiments^8^.

Finally, we combined the tools devised above to investigate how bacterial growth rate modulates the propensities at which lambda enters and exits the lysogenic state. To probe the probability of lysogenization as a function of growth rate, we performed infection at different stages of cell growth (**Figure 4A**) and utilized the lysogen counting method (**Figure 3** above) to measure the frequency of lysogeny at varying MOIs (see ***SI Methods***). The lysogeny-vs.-MOI curve at each growth rate (**Figure 4B**) was then used to infer the single-cell propensity to lysogenize (**Figure S12**; see ***SI Methods***). Our analysis revealed that, upon infection by a single phage, the propensity to lysogenize decreases approximately exponentially with growth rate (**Figure 4C**). This finding is consistent with previous reports of increased lysogenization in stationary^30,32^ and starved cells^29,33^, but places them in the broader context of growth-rate dependent lysogenization—often presumed^34^ but (to the best of our knowledge) not previously shown. Upon co-infection by two phages, the probability of lysogenization in growing cells increases 40–80 fold (**Figure 4D**), suggesting that viral self-counting drives the cell fate choice^10^. However, this feature is absent in stationary cells, where higher MOI does not significantly increase lysogenization (**Figures 4D** and **S13**). Utilizing the inferred single-cell lysogenization curves allowed us to reproduce the experimentally measured “fate diagram”^10^ depicting the population-averaged frequency of lysogeny as a function of MOI and bacterial growth rate (**Figure 4E**).

**Figure 4:**
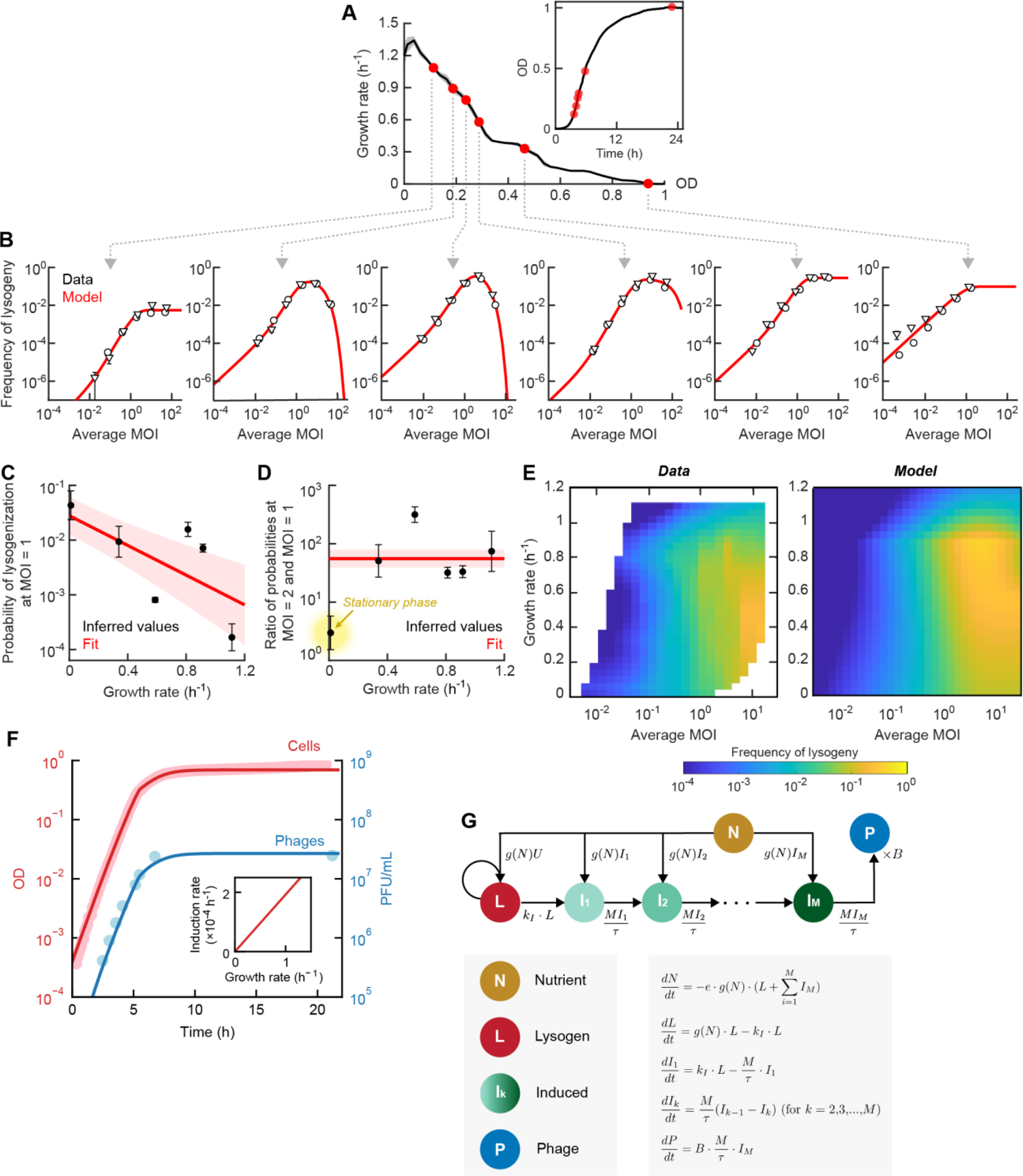
The effect of growth rate on lambda lysogenization and spontaneous induction. **(A)** Performing infection at different growth rates. Black line, the growth rate, as a function of OD, of an uninfected culture of *E. coli* MG1655 grown at 30°C in LBM; gray shading, SEM from triplicate cultures. Red markers, the growth rate at which lambda phages (λ_ts_) were added. Inset, the growth curve of the same culture. **(B)** The frequency of lysogeny as a function of MOI (adjusted for the fraction of phage-infected cells), measured at different bacterial growth rates. Circles and triangles, data obtained in two independent runs of the experiment; error bars, SEM. Red lines, model fits. **(C)** The inferred probability of lysogenization in cells with MOI = 1 as a function of growth rate. Markers, fitted values; error bars, SEM from two independent runs of the experiment. Red line, exponential fit, serving as a guide to the eye. Red shading, fitting uncertainty obtained by bootstrapping. **(D)** The inferred ratio of lysogenization probabilities at MOI = 2 and MOI = 1, as a function of growth rate. Markers, fitted values; error bars, SEM from two independent runs of the experiment. Red line, linear fit, serving as a guide to the eye. Red shading, fitting uncertainty obtained by bootstrapping. Stationary cells (yellow highlight) do not exhibit an increase in the probability of lysogeny between MOI = 1 and MOI = 2. **(E)** The frequency of lysogeny as a function of MOI and growth rate. Left, interpolated experimental data. Right, model prediction. **(F)** The densities of bacteria (red) and free phages (blue) during growth of lysogens. MG1655 λ_ts_ was grown at 30°C in LBM supplemented with 0.2% glucose. Markers, experimental data. Lines, model fits. Inset, the inferred rate of spontaneous induction as a function of growth rate. **(G)** Schematics and equations of the model used for capturing the data in panel F. Circles, species tracked by the model. Arrows, transitions between species. The transition rates are indicated next to the corresponding arrows.

After characterizing the effect of growth rate on the choice to enter lysogeny, we aimed to identify how it impacts the reverse process of spontaneous induction, where lysogenic cells stochastically switch to the lytic pathway^11,34^. To that end, we tracked the growth of lysogenic cells over 24 hours (MG1655 λ_ts_, grown at 30°C in LB supplemented with 10 mM MgSO_4_ and 0.2% glucose, the latter added to inhibit phage adsorption to cells^13,35^). At various time points, corresponding to different bacterial growth rates, phages were extracted ^11^ and enumerated using our phage counting method (**Figure 1** above; see ***SI Methods***). The coupled growth dynamics of lysogenic bacteria and released phages (**Figure 4F**) were interpreted using a mathematical model (**Figure 4G**) analogous to that in **Figure 2** above. Here, we modeled spontaneous induction as a first-order transition from lysogenic cells (*L*) to induced cells (*I*_1_) with rate constant *k*_*I*_. The latent period was modeled as before, with induced cells undergoing several intermediate states (*I*_2_, …, *I*_*M*_) until lysis (see ***SI Methods*** for the full model, and **Table S7** and **Table S8** for parameter values). Using this model, we found that the spontaneous induction rate scales linearly with the growth rate, from approx. 0 in stationary cells to ≈ 2×10^−4^ induction events per hour in early-exponential cells (**Figure 4F**, inset). This finding is consistent with the recent report that the rate of spontaneous SOS activation, the driver of lytic induction^11^, increases with growth rate^36^. The linearity of the induction rate with the growth rate results in a constant switching rate per generation of approx. 1×10^−4^. This value is similar to the estimate by ref. ^11^, but, as in the case of lysogenization above, generalizes it across the entire growth curve of the culture.

Considered together, our measurements of the rates for entering and exiting lysogeny suggest that dormancy is prioritized under conditions of slow growth: During infection, the probability of lysogenization increases in slower - growing cells (**Figure 4C**); once lysogeny is established, slower cell growth results in a lower rate of spontaneous induction into the lytic state (**Figure 4F**). The idea that slow growth promotes lysogeny is part of the accepted narrative for lambda^34^ and other temperate phages^6^, premised on the rationale that slower growing cells would have reduced capacity for a successful lytic reproduction. To the best of our knowledge, the data presented here provides the first quantitative test for this common narrative. In terms of its mechanistic underpinnings, multiple regulatory interactions feed from the signaling molecules encoding information on cellular growth (ppGpp, cAMP), through cellular proteases (FtsH, Lon, RecA) and ribonucleases (RNase III), into the phage decision circuitry^7,37^. Our data reveals that, as long speculated, these myriad regulatory interactions enable the phage to sense and respond to its host’s growth rate, providing yet another example for temperate phages’ ability to process information in order to choose their developmental path optimally^5,38,39^.

In closing, the simple pipeline presented here enables the counting of bacteriophages in unknown samples and the inference of phage encounter rate, latent period, burst size, and frequencies of entering and exiting dormancy. Streamlining the infection protocol necessitated a removal of several steps commonly included, in particular, cell concentration via centrifugation to accelerate phage adsorption, and a temperature upshift to synchronize phage entry^1–3^. Despite these shortcuts, the infection procedure yielded reproducible dynamics, which were interpretable through the use of mathematical modeling as described above. The simplified procedure has the added benefits of reduced perturbation to host physiology and the capacity to systematically scan infection parameters in a high-throughput manner.

The dominant feature in the growth curves of infected cultures was a single peak followed by massive lysis, a feature whose quantitative characteristics were used for the inference of infection parameters. However, more complex dynamics were observed, reproducibly, under certain infection conditions. Whereas some of the additional growth features—the survival of lysogens, and the growth recovery of resistant cells—were addressable by our model, other features remain outside it. Notably, multiple cycles of growth and lysis were observed following infection at high concentrations of lambda, T4, and T5 (**Figure S14**). These repeated cycles suggest a transient cellular state of insusceptibility to phage infection^27,40,41^, and highlight the potential role of population heterogeneity, a subject that merits further investigation. We expect even richer dynamics to emerge under infection scenarios that involve additional phage-host interactions, such as phage-mediated quorum sensing^42–44^, or, conversely, bacterial anti-phage systems^45–47^. We believe that the approach presented here, combining high-throughput infection with modeling-based interpretation, will prove valuable in illuminating such cases.

## ACKNOWLEDGEMENTS

We are grateful to S. Maslov, K. Sneppen, and all members of the Golding lab for their generous advice. Work in the Golding lab is supported by the National Institutes of Health grant R35 GM140709 and by the Alfred P. Sloan Foundation. We gratefully acknowledge the computing resources provided by the Computational and Integrative Biomedical Research Center of Baylor College of Medicine.

## SUPPLEMENTARY INFORMATION

Methods

Figures S1 to S16

Tables S1 to S9

References for Supplementary Information

## Notes

### Competing Interest Statement

The authors have declared no competing interest.

