## Supplementary Information for "Using population dynamics to count bacteriophages and their lysogens"

***This file includes:***

### SUPPLEMENTARY METHODS

#### 1. Bacterial strains and phages

All strains used in this study are listed in **Table S1**. In most experiments involving phage lambda (**Sections 4, 5, 7–9**), we used a temperature-sensitive phage strain, λ *cI*857 *bor*::*kan*$^R$, hereafter $λ_{ts}$, and an isogenic strain without the *cI* mutation, λ $cI_{wt}$ *bor*::*kan*$^R$, hereafter $λ_{wt}$. Phages carrying the *cI*857 allele cannot establish or maintain lysogeny at 37°C or above[1]. We therefore used $λ_{ts}$ at 38°C ± 0.5°C as an obligately lytic variant of lambda, and $λ_{wt}$ as the corresponding wild-type variant. Both phage strains also harbor a kanamycin resistance cassette, which was used to select for lysogenic cells (**Sections 7, 9** and **10**). Other phages (T4, T5, and P1) were used as described in **Section 4** below.

#### 2. Growth media and conditions

##### 2.1. Growth media

The medium used in most experiments was LB (Lennox formulation)[2], comprising (w/v) 1% tryptone (BD Biosciences), 0.5% yeast extract (BD Biosciences), and 0.5% NaCl (Fisher Scientific); the pH was adjusted using 1 mM NaOH (Fisher Scientific). LBM is LB supplemented with 10 mM $MgSO_4$ (Fisher Scientific). LBMM and LBGM are LBM supplemented with 0.2% maltose (Fisher Scientific) or 0.2% glucose (Fisher Scientific)[1]. In experiments involving T5 and P1 phages, LB was supplemented with 1 mM or 5 mM $CaCl_2$ (Fisher Scientific), respectively[3,4].

Other media used in this study are as follows. Tryptone broth (TB) is composed of (w/v) 1% tryptone and 0.8% NaCl, and TBM is TB supplemented with 10 mM $MgSO_4$. Minimal media are based on the M9 minimal salts broth media without a carbon source (Teknova). M9Mal and M9Glu are M9 broth supplemented with 0.4% maltose or 0.4% glucose, respectively. For the plaque formation assay[5,6], NZYM medium was prepared using (w/v) 2.2% NZYM powder (Teknova); the pH was adjusted using 10 mM NaOH.

Agar plates and soft agar were prepared by supplementing the liquid medium with 1.5% and 0.7% w/v agar (BD Biosciences), respectively[5].

##### 2.2. Growth conditions

To prepare overnight cell cultures, fresh colonies on LB agar plates (supplemented with 50 µg/mL kanamycin, Fisher Scientific, for lysogenic strains) were inoculated into 2 mL of medium (as specified for each experiment below) in 14 mL round-bottom test tubes (Falcon). Overnight cultures were grown for approx. 16 hours at 30°C with aeration (220 rpm shaking). The overnight cultures were diluted into experimental ("overday") culture as described below (**Sections 4**, **5**, **8**, and **9**).

##### 2.3. Measuring the optical density

Throughout this work, we used two different instruments to measure the optical density (OD): (i) When growing cells in bulk cultures, a SmartSpec Plus spectrophotometer (Bio-Rad) was used to measure the OD at a wavelength of 600 nm and a path length of 1 cm. The corresponding values are denoted below as $OD_{spec}$. (ii) During growth in the plate reader, the instrument (TECAN Infinite F200 Pro and TECAN M Nano) measured the OD at 595 nm and a path length determined by the depth of liquid in the well (≈ 3 mm in our experiments). We denoted these values, used in the analysis of growth dynamics, simply as "OD". The two measurements differ by a scaling factor, $OD_{spec} \div OD = 4.51 \pm 0.05$.

### 3. Phage preparation

#### 3.1. Preparing lambda phages

We followed the protocols in refs. [1,7] to produce crude phage lysates. For the temperature-sensitive $\lambda_{ts}$, we performed a heat induction of the lysogens. Briefly, an overnight culture of MG1655 $\lambda_{ts}$ was diluted 1000-fold into LBGM in a baffled Erlenmeyer flask, and incubated at 30°C with mild shaking (180 rpm). Upon reaching $OD_{spec} \approx 0.4$, the culture was incubated in a water bath at 42°C with 180 rpm shaking for 15 minutes, then at 37°C with 180 rpm shaking for 1 hour until $OD_{spec}$ dropped below 0.05. For $\lambda_{wt}$, we performed a chemical induction using mitomycin C (MMC, FisherScientific). Briefly, an overnight culture of MG1655 $\lambda_{wt}$ was diluted 1000-fold into LBGM in a baffled Erlenmeyer flask, and incubated at 37°C with mild shaking (180 rpm). At $OD_{spec} \approx 0.4$, 10 µg/mL of MMC was added to the culture. The flask was wrapped in foil, and incubation (37°C, 180 rpm shaking) was resumed for 2–3 hours. Lysis was determined to be complete when the $OD_{spec}$ dropped below 0.2.

Following either heat or MMC induction, the lysed culture was supplemented with 5% chloroform, and incubated at room temperature (RT) for 15 minutes. The lysate was centrifuged at 4000×g for 10 minutes at 4°C to pellet the debris, and the clear supernatant was extracted, supplemented with 0.3% chloroform, and stored at 4°C until use. Standard plaque formation assays were performed, using NZYM agar, to determine the phage titers ($\approx 10^{10}$ plaque-forming units, PFU/mL).

When higher titers were required, we used the crude induction lysates (produced using heat induction) to perform phage precipitation using polyethylene glycol (PEG), followed by resuspension of the phage pellets in SM buffer (Teknova) as described in ref. [8]. These additional steps increased the phage titer to $\approx 10^{11}$ PFU/mL.

#### 3.2. Preparing other phages

Phages T4, T5, and P1 were produced by infecting cell cultures, as described in ref. [2]. Briefly, cultures of MG1655 were grown at 37°C in LB (supplemented with 1 mM $CaCl_2$ for T5, as described in ref. [3], or 5 mM $CaCl_2$ for P1, as described in ref. [4]). When the culture reached $OD_{spec} \approx 0.2$, approximately $10^7$ PFUs of T4, T5, or P1 were added to the cultures. The infected cultures were incubated at 37°C without shaking for 15 minutes, then shaking was resumed until lysis was observed (approximately 3–4 hours). Chloroform treatment and centrifugation were then performed as described in **Section 3.1** above.

### 4. Microplate-based infection assay for measuring phage concentrations

In **Sections 4.1–4.3** below, we describe the assay for infection by $\lambda_{ts}$ in LBM. Assays involving other phages and other media are described in **Section 4.4**.

#### 4.1. Calibration curves using known phage concentrations

##### *4.1.1. Preparing the phage standards*

A 10-fold serial dilution of $\lambda_{ts}$ phages (prepared as described in **Section 3**), with concentrations spanning from $6\times10^2$ PFU/mL to $6\times10^{11}$ PFU/mL, was prepared in SM buffer.

##### *4.1.2. Cell cultures and infection*

An overnight culture of MG1655 in LB (prepared as described in **Section 2**) was diluted 1:500 into LBM in a baffled Erlenmeyer flask. The culture was then grown at 37°C with 220 rpm shaking. Upon reaching $OD_{spec} \approx 0.1$, the culture was diluted 1:10 into LBM and aliquoted into a clear 48-well flat-bottom microplate (COSTAR), with each well containing 500 µL of culture. As negative controls for cell growth, some wells containing blank LBM were included in the microplate. The microplate was then placed into the plate reader, incubated at 38°C with shaking (orbital mode, 1 mm amplitude), and the OD was recorded every 5 minutes.

After 30 minutes of growth in the plate reader, 10 µL of phages at different concentrations were added to each well. We included two replicates for each phage concentration. As negative controls for phage infection, we included two uninfected cultures to which blank SM buffer was added. After phage addition, incubation was resumed for at least 12 hours.

*4.1.3. Data analysis*

The average OD in the wells with blank LBM was first subtracted from all OD measurements. Next, we identified the first local maximum in the growth curve for each culture, which, for infected cultures, corresponds to the onset of massive lysis. We term this OD value the “lysis OD”. For lambda phages infecting cells in LBM medium, the relationship between the lysis OD and the logarithm of the initial phage concentration is approximately linear (**Figure 1B**). The calibration curve was obtained by fitting a linear equation:

$$y = k \cdot x + b, \tag{4.1}$$

where $y$ is the lysis OD and $x$ is the logarithm of the initial phage concentration in PFU/mL.

##### 4.2. Measuring phage concentrations in unknown samples

For a sample with unknown phage concentration, the same infection protocol was performed as described in **Section 4.1.2**. The lysis OD was identified using the same procedure in **Section 4.1.3**. The calibration curve (**Equation 4.1**) was used to calculate the unknown phage concentration corresponding to this lysis OD.

##### 4.3. Measuring phage concentrations in samples extracted from infected cultures

To measure the phage concentrations in the infected cultures during cell growth (**Figure 2D**), we extracted phages as follows. At each time point (before phage addition, then at 1, 2, 3, 4, 5, 6, 7, 8, and 20 hours after phage addition), 5 μL of the infected cultures were taken from the wells and diluted into 495 μL of LBM media (a 1:100 dilution). Then, 25 μL of chloroform (final concentration, 5%) was added to each culture, followed by vortexing for 10 seconds, to lyse the cells[1]. Phage lysates were stored at 4°C, and the phage counting procedure (**Section 4.2**) was used to measure the phage concentrations in each sample.

##### 4.4. Calibration curves for infection in other media or by other phages

We also performed the calibration assay (**Section 4.1**) for infection in other growth media, other lambda strains, or other phages. For infections in complex media (LB, LB supplemented with $CaCl_2$, and TBM), the calibration curves were analyzed using **Equation 4.1**. For infections in minimal media (M9Glu and M9Mal), the lysis OD was found to be approximately a power function of the initial phage concentration. The calibration curve was obtained by fitting the following equation:

$$\log_{10} y = k \cdot x + b, \tag{4.2}$$

where $y$ and $x$ are defined the same way as in **Equation 4.1**. All calibration curves under different infection conditions are shown in **Figure S3.**

#### 5. Detecting single phages

##### 5.1. Phage preparation

A solution of $\lambda_{ts}$ (prepared as described in **Section 3**) was diluted in SM to reach a concentration of $\approx 10^4$ PFU/mL. The concentration was confirmed using a plaque assay on NZYM agar plates[5]. Immediately before infection, the phage solution was further diluted 1:200 in SM (thus reaching 38 PFU/mL, or 0.38 PFU in 10 μL on average).

#### 5.2. Cell cultures and infection

Cells were cultured in LBMM and aliquoted into a 48-well microplate as described in **Section 4.1**. After 30 minutes of growth in the plate reader, 10 µL of phage solution (containing, on average, 0.38 PFU) were added to each culture well. We included 32 replicates of cultures with phages and 10 replicates without phages. After phage addition, incubation was resumed, and the OD was recorded for 48 hours.

When the same assay was performed in LBM, we noticed that the fraction of lysed cultures (analyzed as described in **Section 5.3** below) was lower than our theoretical expectation. We reasoned that the cultures in LBM had entered the stationary phase before massive lysis occurred. To extend the range of OD over which the cultures can grow before saturation, we therefore used LBMM in this assay instead.

#### 5.3. Data analysis

##### *5.3.1. Predicting the expected number of lysed cultures*

We denoted the average number of phage particles in 10 µL as $\lambda$ (dictated by the experimental design). The number of phage particles in each of the 10 µL aliquots ($X$) is expected to follow the Poisson distribution:

$$P(X = x) = \frac{\lambda^x e^{-\lambda}}{x!}. \quad (5.1)$$

The probability that there is a non-zero number of phage particles in that aliquot volume, $\pi_0$, is:

$$\pi_0 = P(X > 0) = 1 - P(X = 0) = 1 - e^{-\lambda}. \quad (5.2)$$

With $\lambda = 0.38$, $\pi_0 = 0.316$.

Among $N$ independent cultures, the number of cultures containing a non-zero number of phage particles ($Y$) follows the Binomial distribution:

$$P(Y = y) = \binom{N}{y} \pi_0^y (1 - \pi_0)^{N-y}. \quad (5.3)$$

Using **Equation 5.3**, we calculated the expected number of $Y$ in $N = 32$ independent cultures. The expected fraction of lysed cultures, $Y/N$, is shown in **Figure 1C, right**.

##### *5.3.2 Identifying lysed cultures*

Massive lysis was identified as described in **Section 4.1.3**. We note that for uninfected cells over 48 hours of growth, the OD displayed a minor decline during the stationary phase, presumably due to cell death[9]. However, this decline is not as drastic as that in massive lysis (**Figure 1C, left**). Therefore, to quantitatively distinguish massive lysis from the death phase, we calculated the difference in OD between the first local maximum and the subsequent minimum for each culture (denoted ΔOD). For the infected samples, the values of ΔOD fell into two distinct groups (**Figure 1C, middle**). One group (ΔOD < 0.3, arbitrarily chosen) was similar to the uninfected cultures, and was classified as unlysed. The other (ΔOD ≥ 0.3) was recorded as lysed. The measured fraction of lysed cultures (10 out of 32 cultures) is shown in **Figure 1C, right**.

##### *5.3.3. Performing a binomial test*

We defined $\pi_0$ as in **Equation 5.2**, and $\pi$ as the observed probability that the 10 µL phage aliquot leads to massive lysis. If a single phage leads to massive lysis, $\pi$ should be equal to $\pi_0$.

We performed a two-tailed binomial test[10] to test the following hypothesis: $H_0 : \pi = \pi_0$. Denoting the observed number of lysis events in $N$ infected cultures as $k$, we calculated the two-tailed *p*-value as follows:

$$p = \sum_{i \in \mathcal{J}} P(Y = i) = \sum_{i \in \mathcal{J}} \binom{N}{i} \pi_0^i (1 - \pi_0)^{N-i}, \tag{5.4}$$

where $\mathcal{J}$ was defined as $\mathcal{J} = \{i: P(Y = i) < P(Y = k)\}$. With $N = 32$ and $k = 10$, the calculated $p$-value was 0.849; as a result, we accepted the null hypothesis and concluded that our assay can detect single phages at the expected efficiency.

### 6. Modeling the growth dynamics of phages and bacteria during infection

##### 6.1. Model description

We aimed to capture the population dynamics, up to but excluding the recovery of bacterial growth (**Figure 1A**). Following refs. [11,12], we used a set of ordinary differential equations (ODE) to describe the dynamics of nutrient resources ($N$), uninfected cells ($U$), infected cells ($I$), and free phages ($P$). The assumptions of the model, shown in **Figure 2A**, are summarized below.

(1) Cell growth: As in ref. [13], we assumed the instantaneous growth rate of uninfected cells $g(N)$ depends on the nutrients ($N$), in the following manner:

$$g(N) = v \cdot \frac{N}{N + K}, \tag{6.1}$$

where $K$ is the Monod constant and $v$ is the maximal growth rate under a given nutrient.

(2) Phage-cell encounter: We assumed that phages ($P$) and cells ($U$ and $I$) encounter each other with a second-order rate constant $r$. The encounter of phages and uninfected cells ($U$) results in the production of infected cells ($I$) (ref. [12]).

(3) Nutrient consumption: The infected cells ($I$) are assumed to consume nutrient resources even if they do not grow. Therefore, the rate of resource consumption is proportional to the combined densities of uninfected and infected cells, the instantaneous growth rate $g(N)$, and a conversion efficiency parameter $e$ that relates cell growth to nutrient consumption[12].

(4) Cell lysis: We assumed the infected cells go through $M$ intermediate states ($I_1, I_2, \ldots, I_M$) before lysis, and the transition rates from one state to the next are identical ($M/\tau$) (refs. [14,15]). The exit from the last state ($I_M$) leads to cell lysis. Therefore, the time between infection and cell lysis (the latent period) follows an Erlang distribution with mean $\tau$ and shape parameter $M$. The larger $M$ is, the narrower the latent period distribution.

(5) Phage release: The number of phages released upon cell lysis (burst size[16]) is denoted as $B$.

Taken together, the population dynamics of nutrient resources, bacterial cells, and free phages obey the following equations (**Equation 6.2**):

$$
\begin{aligned}
&\frac{dN}{dt} = -e \cdot \left(U + \sum_{i=1}^{M} I_i\right) \cdot g(N),\\
&\frac{dU}{dt} = U \cdot g(N) - r \cdot U \cdot P,\\
&\frac{dI_1}{dt} = r \cdot U \cdot P - \frac{M}{\tau} \cdot I_1,\\
&\frac{dI_i}{dt} = \frac{M}{\tau} \cdot (I_{i-1} - I_i) \text{ for } i = 2,3,\dots,M,\\
&\frac{dP}{dt} = B \cdot \frac{M}{\tau} \cdot I_M - r \cdot \left(U + \sum_{i=1}^{M} I_i\right) \cdot P.
\end{aligned}
\tag{6.2}
$$

To fit this model and its variations (described in **Sections 6.6, 6.7, and 8.2**), the initial nutrient concentration $N(0)$ was set to be 1, and the cell concentrations were divided by $10^9$ colony-forming units, CFU/mL, which corresponds to the approximate cell concentration at OD = 1.

##### 6.2. Parameterization of the cell growth

We assumed nutrients are consumed sequentially[17,18] through $X$ phases. In each phase $(i)$, cell growth is controlled by one limiting substrate via substrate-specific $v_i$ and $K_i$, where $i = 1,2,\dots,X$. The transition between these phases is defined by the thresholds $\theta_i$, where $\theta_1 = 1$ (the first growth phase with maximum nutrient), and $\theta_{X+1}$ (the final growth phase) $= 0$. When $N$ decreases below $\theta_{i+1}$, the substrate $i$ is considered exhausted, and cells begin to consume substrate $i + 1$. We also assumed that the conversion factor from cell growth to nutrient consumption is $e$. Therefore, the dynamics of nutrients and cells in the absence of infection are described by the following equations (**Equation 6.3**):

$$
\begin{aligned}
&\text{When } \theta_{i+1} \le N < \theta_i,\\
&\frac{dN}{dt} = -e \cdot U \cdot v_i \cdot \frac{N}{N + K_i},\\
&\frac{dU}{dt} = U \cdot v_i \cdot \frac{N}{N + K_i}.
\end{aligned}
\tag{6.3}
$$

We scanned $X$ from 1 to 3, and fitted the model above to the OD curves of uninfected cultures by minimizing the following objective function:

$$
R(\{v_i\},\{K_i\},\{N_i\}) = \sum_{t \in T_0} |OD(t) - U(t)|,
\tag{6.4}
$$

where $OD(t)$ is the measured OD of uninfected cultures at time $t$, $U(t)$ is the model-predicted density of uninfected cells, and $T_0$ is the set of time points where OD was measured. **Equation 6.4** was minimized using the simulated annealing algorithm[19].

Following this analysis, we found that growth in single-substrate media (M9Glu and M9Mal) was describable using $X = 1$, with a single set of $v$ and $K$ (see **Figures S4C** and **S4D** and **Table S2**). Growth in complex media (e.g., LBM and TBM), was captured by models with multiple growth phases ($X = 2$ for TBM and $X = 3$ for LBM, **Figures S4B** and **2B**, **Tables S3** and **S4**).

##### 6.3. Parameterization of the latent period

Considering the growth dynamics during the interval between massive lysis and onset of growth recovery (**Figure 1A**), and assuming that the contribution of uninfected cells to OD is negligible, we wrote:

$$\frac{dI_1}{dt} = -\frac{M}{\tau} \cdot I_1,$$
$$\frac{dI_i}{dt} = \frac{M}{\tau} \cdot (I_{i-1} - I_i) \text{ for } i = 2,3,\ldots,M, \tag{6.5}$$

where $I_i$ describes the different states of the infected cells between phage-cell encounter and cell lysis ($i = 1,2,\ldots,M$).

Following refs. [14,15], we scanned $M$ from 1 to 10, and fitted the mean latent period $\tau_j$ for each infection condition $j$ (defined by the initial phage concentration, $P_j(0)$). Specifically, we minimized the following objective function, using simulated annealing:

$$R(\tau_{j,M}) = \sum_{t \in T_o} \left| OD_j(t) - \sum_{i=1}^{M} I_{i,j}(t) \right|, \tag{6.6}$$

where $OD_j(t)$ is the measured OD dynamics in condition $j$, $T_o$ is the set of time points where OD concentrations were measured, and $I_{i,j}$ is the predicted $I_i$ for infection condition $j$. This procedure gives a set of best-fitted $\{\tau_{j,M}\}$ for a given $M$. Using these sets of $\{\tau_{j,M}\}$, we calculated the mean absolute error (MAE) for each $M$:

$$\mathrm{MAE}(M) = \sum_j \sum_{t \in T_o} \left| OD_j(t) - \sum_{i=1}^{M} I_{i,j}(t) \right|. \tag{6.7}$$

For infection by $\lambda_{ts}$ in LBM, the OD curves of cultures infected at six different phage concentrations ($1.2\times10^8$, $1.2\times10^7$, $1.2\times10^6$, $1.2\times10^5$, $1.2\times10^4$, and $1.2\times10^3$ PFU/mL) were used for fitting. The MAE was minimized when $M = 5$ (**Figure 2C, left**). Therefore, we chose $M = 5$ and the associated $\{\tau_{j,5}\}$ in the following analysis (**Section 6.4**).

For infection in other media and infection by other phages, a similar fitting strategy was applied. The values of $M$ under different infection conditions are shown in **Table S5**.

##### 6.4. Characterizing latent period as a function of growth rate

We further parameterized the latent period $\tau$ (obtained as described in from **Section 6.3**) as a function of the instantaneous growth rate $g$.

We defined the instantaneous growth rate at lysis OD as the growth rate of the uninfected culture at the same OD. The instantaneous growth rate was obtained by fitting $\log OD(t)$ as a linear function of time, with the previous and next two time points included (i.e., over 20 minutes). Examining the data, we found that the latent period ($\tau$) is well-approximated by a linear function of the inverse lysis-OD growth rate ($1/g$), shown in **Figure 2C, right**:

$$\tau = b + k \cdot \frac{1}{g}. \tag{6.8}$$

Therefore, we used **Equation 6.8** to predict the latent period at a given growth rate, $\tau(g)$. For infection in other media and infection by other phages, a similar fitting strategy was applied. The fitted parameter values are shown in **Table S5**.

##### 6.5. Parameterization of phage/cell encounter rate and burst size

After cell growth and latent period have been parameterized, the only remaining parameters in **Equation 6.2** are the phage/cell encounter rate $r$ and the burst size $B$. To estimate these parameters, we fitted the model to the OD curves of cultures infected at six different initial phage concentrations and to the concentrations of phages extracted from the same cultures. The procedure for determining the phage concentrations is described in **Section 4.3**.

The fitting was performed by minimizing the objective function:

$$R(r,B) = \sum_{j=1}^{6}\left[\frac{1}{|T_o|}\sum_{t\in T_{o,j}}\frac{\left|\left(U_j(t)+\sum_{i=1}^{M} I_{i,j}(t)\right)-OD_j(t)\right|}{\left|\max OD_j(t)-\min OD_j(t)\right|} + \frac{1}{|T_P|}\sum_{t\in T_P}\frac{\left|\log \widehat{P}_j(t)-\log P_j(t)\right|}{\left|\max\left(\log P_j(t)\right)-\min\left(\log P_j(t)\right)\right|}\right], \tag{6.9}$$

where $j$ indexes the infection conditions, $\widehat{P}_j(t)$ is the model-predicted phage dynamics for infection condition $j$, and $OD_j(t)$ and $P_j(t)$ are the measured OD and phage dynamics. $T_{o,j}$ is the set of time points where the measured OD concentrations were used for fitting. $T_P$ are the set of time points where phage concentrations were measured. $|T_{o,j}|$ and $|T_P|$ are the corresponding total numbers of time points.

For infection by $\lambda_{ts}$ in LBM, all measured phage concentrations were used in fitting, while only the OD data points whose second-order time-derivative $\frac{d^2OD}{dt^2}$ (approximated using the centered finite differences) is greater than $3.5\times10^{-5}$ were used in fitting. The fitting results are shown in **Figure 2D**, and the parameter values are listed in **Table S6**.

For infection by other phages and in other growth media, since phage concentrations were not measured, we only used the OD data for fitting. The fitting results are shown in **Figure S6** and the fitted parameters are listed in **Table S6**.

##### 6.6. Modeling bacterial recovery

We aimed to incorporate the recovery after massive lysis into the framework developed in **Section 6.1**. Following ref. [12], we assumed that three processes contribute to this recovery:

(1) Growth of cells that are resistant to phage infection;

(2) Conversion from lysed cells to debris, which contributes to the measured OD;

(3) Recycling of nutrients from cell debris into the nutrients, which foster cell growth.

We denoted the resistant population as $R$ and cell debris as $D$ (**Figure S7A**). We assumed that the resistant cells were produced from uninfected cells ($U$) with a first-order transition rate $k_m$ (refs. [11,12]), and all lysed cells were converted to cell debris instantaneously. The content of each lysed cell was recycled as nutrients ($N$) with a conversion factor $k_f$. Finally, we assumed that the debris from each lysed cell contributed to OD with a 10% extinction coefficient of an intact cell (a value inferred from the OD curves of heat-induced temperature-sensitive lysogens, and consistent with ref. [20]).

Accordingly, we modified **Equation 6.2** to describe the dynamics of nutrients ($N$), uninfected cells ($U$), infected cells ($I_i$), free phages ($P$), resistant cells ($R$) and cell debris ($D$):

$$
\begin{aligned}
\frac{dN}{dt} &= -e \cdot \left(U + R + \sum_{i=1}^{M} I_i\right) \cdot g(N) + k_f \cdot \frac{M}{\tau} \cdot I_M, \\
\frac{dU}{dt} &= U \cdot g(N) - r \cdot U \cdot P - k_m \cdot U, \\
\frac{dI_1}{dt} &= r \cdot U \cdot P - \frac{M}{\tau} \cdot I_1, \\
\frac{dI_i}{dt} &= \frac{M}{\tau} \cdot (I_{i-1} - I_i) \text{ for } i = 2,3,\dots,M, \\
\frac{dP}{dt} &= B \cdot \frac{M}{\tau} \cdot I_M - r \cdot \left(U + \sum_{i=1}^{M} I_i\right) \cdot P, \\
\frac{dR}{dt} &= R \cdot g(N) + k_m \cdot U, \\
\frac{dD}{dt} &= \frac{M}{\tau} \cdot I_M.
\end{aligned}
\tag{6.10}
$$

Since $g$ was parameterized from the OD dynamics of uninfected cultures, and $\tau$, $r$, and $B$ were parameterized from early-stage infection data (as described in **Sections 6.1–6.5**), the only remaining unknown parameters are $k_m$ and $k_f$. We fitted $k_m$ and $k_f$ by minimizing the following objective function:

$$
\begin{aligned}
R(r,B) = \sum_{j=1}^{6} \Bigg[ & \frac{1}{|T_{o,j}|} \sum_{t \in T_{o,j}} \frac{\left|\left(U_j(t) + R_j(t) + 0.1 \cdot D_j(t) + \sum_{i=1}^{M} I_{i,j}(t)\right) - OD_j(t)\right|}{\left|\max OD_j(t) - \min OD_j(t)\right|} \\
& + \frac{1}{|T_P|} \sum_{t \in T_P} \frac{\left|\log \widehat{P}_j(t) - \log P_j(t)\right|}{\left|\max\left(\log P_j(t)\right) - \min\left(\log P_j(t)\right)\right|} \Bigg],
\end{aligned}
\tag{6.11}
$$

using the same notation as shown in **Equation 6.9**.

##### 6.7. Modeling temperate phage infection

To describe the population dynamics after infection by temperate phages, we extended the framework developed in **Section 6.6** by adding a lysogenic population ($L$), produced with a constant frequency $q$ at each infection event (**Figure S9A**).

The following equations describe the dynamics of nutrients ($N$), uninfected cells ($U$), infected cells ($I_i$), free phages ($P$), resistant cells ($R$), cell debris ($D$) and lysogens ($L$):

$$
\begin{aligned}
&\frac{dN}{dt} = -e \cdot \left( U + R + L + \sum_{i=1}^{M} I_i \right) \cdot g(N) + k_f \cdot \frac{M}{\tau} \cdot I_M, \\
&\frac{dU}{dt} = U \cdot g(N) - r \cdot U \cdot P - k_m \cdot U, \\
&\frac{dI_1}{dt} = r \cdot U \cdot P - \frac{M}{\tau} \cdot I_1, \\
&\frac{dI_i}{dt} = \frac{M}{\tau} \cdot (I_{i-1} - I_i) \text{ for } i = 2,3,\dots,M, \\
&\frac{dP}{dt} = B \cdot \frac{M}{\tau} \cdot I_M \cdot (1-q) - r \cdot \left( U + L + \sum_{i=1}^{M} I_i \right) \cdot P, \\
&\frac{dR}{dt} = R \cdot g(N) + k_m \cdot U, \\
&\frac{dD}{dt} = \frac{M}{\tau} \cdot I_M \cdot (1-q), \\
&\frac{dL}{dt} = \frac{M}{\tau} \cdot I_M \cdot q.
\end{aligned}
\tag{6.12}
$$

All parameters except $k_m$, $k_f$, and $q$ were parameterized as described in **Sections 6.1–6.5**. $k_m$ and $k_f$ were estimated from the infection data by virulent phages (**Section 6.6**). Therefore, the only remaining unknown parameter is $q$.

We fitted the frequency of lysogeny, $q_j$, for each infection condition $j$, by minimizing the following objective function:

$$
R(q_j) = \frac{1}{|T_{o,j}|} \sum_{t \in T_{o,j}} \frac{\left| \left( U_j(t) + R_j(t) + L_j(t) + 0.1 \cdot D_j(t) + \sum_{i=1}^{M} I_{i,j}(t) \right) - OD_j(t) \right|}{\left| \max OD_j(t) - \min OD_j(t) \right|}, \tag{6.13}
$$

using the same notation as shown in **Equation 6.9**. The fitting results for infection by $\lambda_{wt}$ in LBM are shown in **Figure S9B**, and the values of $q_j$ are shown in **Figure S9C**.

### 7. Quantifying the proportion of lysogens among surviving cells in phage-infected cultures

To verify that the surviving cells in cultures infected by $\lambda_{wt}$ (**Section 4**) were lysogens, we leveraged the fact that $\lambda_{wt}$ harbors a kanamycin resistance cassette[1]. After the cell culture infected at an initial phage concentration of ≈ $2\times10^7$ PFU/mL had exhibited massive lysis, the culture was extracted and diluted $4\times10^4$-fold using 1× PBS. Diluted cells were plated on agar plates made using LB or LB supplemented with 50 μg/mL kanamycin. The numbers of colonies were used to calculate the total number of cells in the infected culture and the number of lysogenic cells (resistant to kanamycin). The results, shown in **Figure S8**, indicated that >99% of the surviving cells were lysogens.

### 8. Measuring the spontaneous induction rate of lambda lysogens as a function of growth rate

##### 8.1. Measuring the number of phages released by lambda lysogens at different growth rates

This assay is modified from ref. [1]. Briefly, an overnight culture of MG1655 $\lambda_{ts}$ in LB, supplemented with 50 µg/mL kanamycin (prepared as described in **Section 2**), was centrifuged, and the supernatant (containing free phages released during overnight growth) was removed. The cell pellet was resuspended in the same volume of fresh LBGM, and further diluted 1000-fold in LBGM. 500 µL of this diluted culture was aliquoted into replicate wells in a clear 48-well flat-bottom microplate (COSTAR). The plate was incubated for 24 hours at 30°C with shaking.

We sampled the bacterial cultures when they were first inoculated, and when the blank-subtracted OD reached approximately 0.01, 0.02, 0.04, 0.25, 0.30, 0.50, and 1.00. At each time point, the entire 500 µL of the cultures from two wells were extracted, and 25 µL of chloroform (final concentration, 5%) was added to each sample, followed by vortexing for 10 seconds. Phage lysates were stored at 4°C, and the phage counting procedure (**Section 4.2**) was used to measure the phage concentrations, with the calibration curve obtained by infection of the same phage strain in LBM (**Figure 1B**).

##### 8.2. Modeling spontaneous induction

We assumed the lysogenic cells $(L)$ switch to the induced state $(I_i)$ with a first-order transition rate $k_I$ (the spontaneous induction rate). The induced cells undergo $M$ intermediate states $(I_1, I_2, \dots, I_M)$ before reaching lysis, similar to the infected cells described in **Section 6.1**. The model schematics are shown in **Figure 4G**.

Accordingly, we modified **Equation 6.2** to describe the dynamics of nutrients $(N)$, lysogens $(L)$, induced cells $(I_i)$ and free phages $(P)$.

$$\begin{aligned}
\frac{dN}{dt} &= -e \cdot \Big(L + \sum_{i=1}^{M} I_i\Big) \cdot g(N), \\
\frac{dL}{dt} &= L \cdot g(N) - k_I \cdot L, \\
\frac{dI_1}{dt} &= k_I \cdot L - \frac{M}{\tau} \cdot I_1, \\
\frac{dI_i}{dt} &= \frac{M}{\tau} \cdot (I_{i-1} - I_i) \text{ for } i = 2,3,\dots,M, \\
\frac{dP}{dt} &= B \cdot \frac{M}{\tau} \cdot I_M.
\end{aligned} \tag{8.1}$$

To parameterize the growth rate, the OD curves of overday cultures in LBGM in the first 12 hours were fitted to **Equation 6.3**. The fitted parameter values are listed in **Table S7**. The data was well described by three growth phases $(X = 3)$ (**Figure S4A**).

The remaining parameters ($k_I$ and $B$) were fitted to the measured dynamics of OD and phage concentration, by minimizing the objective function:

$$R(k_I, B) = \frac{1}{|T_o|} \sum_{t \in T_o} \frac{\left|\left(L(t) + \sum_{i=1}^{M} I_i(t)\right) - OD(t)\right|}{|\max OD(t) - \min OD(t)|} + \frac{1}{|T_P|} \sum_{t \in T_P} \frac{\left|\log \hat{P}(t) - \log P(t)\right|}{|\max(\log P(t)) - \min(\log P(t))|}, \tag{8.2}$$

using the same notation as shown in **Equation 6.9**. The fitting result (**Figure S15**) shows that this model overestimated the phage concentration at slow bacterial growth.

To better capture the data, we incorporated into the model a dependence of the spontaneous induction rate $k_I$ on the normalized bacterial growth rate $\phi(N)$:

$$\phi(N) = \frac{g(N)}{\max g(N)}, \tag{8.3}$$

where $\max g(N)$ was obtained when $N = 1$. Specifically, we assumed that $k_I$ is a linear function of $\phi(N)$:

$$k_I = k \cdot \phi(N) + b. \tag{8.4}$$

The fitting results are shown in **Figure 4F**, and the fitting parameters are provided in **Table S8**.

#### 9. Measuring the frequency of lysogeny as a function of MOI and growth rate

##### 9.1. Infecting at different MOIs at a given growth rate

For this assay, we adapted the bulk lysogenization protocol from refs. [5,21], but instead of plating for colonies to measure the concentration of cells, we used the growth dynamics of the infected bacterial cultures as described below. This assay was used to produce the data shown in **Figures 3B** and **3C**.

###### *9.1.1. Preparing phages*

A 2-fold dilution series of $\lambda_{ts}$ (harboring a kanamycin resistance cassette, produced as described in **Sections 3**) was prepared in LBM. The concentrations of phages in this dilution series ranged from ≈ $5\times10^7$ to ≈ $2\times10^{11}$ PFU/mL.

###### *9.1.2. Cell culturing and infection*

To prepare cells, an overnight culture of MG1655 in LB (prepared as described in **Section 2**) was diluted 1:500 into LBM in a baffled Erlenmeyer flask. This culture was grown at 30ºC with aeration (220 rpm shaking). Upon reaching $OD_{spec}$ ≈ 0.1, 500 µL of the culture was aliquoted into different wells of a clear 48-well flat-bottom microplate (COSTAR). This plate (“infection plate”) was incubated for 30 minutes at 30ºC with shaking in a plate reader, where the optical density (OD) was recorded every 5 minutes.

After 30 minutes, 10 µL of phages at different concentrations were added to different wells, resulting in infected bacterial cultures at multiplicity of infection, MOI, ranging from ~0.02 to ~100. As negative controls for phage infection, we included uninfected cultures, to which blank SM buffer was added. The infection plate was incubated with shaking at 30ºC for 15 minutes to allow phages to infect cells. Then, 2 µL of each sample was diluted into 500 µL of LBM in another 48-well microplate (“detection plate”), pre-warmed at 30ºC. The detection plate was incubated with shaking at 30ºC for 45 minutes. Then, each sample was supplemented with kanamycin (final concentration, 50 µg/mL), and incubated for 24 hours.

For the uninfected samples, some wells were supplemented with 50 µg/mL kanamycin, serving as a negative control for infection, whereas some were not subjected to selection, providing an estimate for the total density of cells in the infection mixture (analyzed as described below).

##### 9.2. Infecting at different MOIs and growth rates

This assay was used to produce the data shown in **Figure 4B**.

*9.2.1. Preparing cells at different growth rates*

To obtain cells at different growth rates, we first prepared a culture of MG1655 in LBM at 30°C, as described in **Section 9.1.2**. At $OD_{spec} \approx 0.1$, this culture was diluted into fresh LBM in different baffled Erlenmeyer flasks (5 cultures, dilution ratios ranging from 1:50 to undiluted). These cultures with different initial OD were grown at 30°C for 3 hours (final $OD_{spec}$ ranging from ~0.2 to ~2.5). Then, 200 µL of these cultures and the overnight culture ($OD_{spec} \approx 5$), were aliquoted into different wells of a 96-well "infection plate", yielding 6 infection series at different growth rates. These cultures were grown for 30 minutes at 30°C before phage addition.

*9.2.2. Infection*

Infection was performed as described in **Section 9.1.2.**, using phages in a 5× dilution series (6 levels, ranging from $\approx 8\times10^7$ to $\approx 2\times10^{11}$ PFU/mL). The infected samples were grown for 30 minutes; then, 1 µL of each sample in the infection plate was diluted into 200 µL of pre-warmed LBM supplemented with 50 µg/mL kanamycin in another 96-well microplate ("detection plate"). The detection plate was incubated with shaking at 30°C for 24 hours. The detection plate also contained the uninfected control cultures with and without kanamycin selection, as described in **Section 9.1.2**.

In contrast to **Section 9.1**, here, we introduced kanamycin selection immediately after the dilution step. Growth in fresh medium without kanamycin selection was omitted to ensure that the density of lysogens we measured reflected lysogenization at the original growth rates.

##### 9.3. Data analysis

*9.3.1. Estimating the growth rate at which infection was performed*

The growth rate at which infection was performed, $g$, was calculated by fitting the following equation to the growth curves of each sample during the 30-minute duration before phages were added.

$$N(t) = N_0 \times e^{g \cdot t}, \tag{9.1}$$

where $N_0$ is the initial cell concentration, and $g$ is the growth rate. Fitting was performed in logarithmic space.

*9.3.2. Calculating the frequency of lysogeny*

Following previous studies[5,21], we defined the frequency of lysogeny as the fraction of kanamycin-resistant lysogenic cells ($L_0$) among all cells in the infected cultures ($T_0$):

$$f_{\text{lysogeny}} = \frac{L_0}{T_0}. \tag{9.2}$$

We inferred $L_0$ and $T_0$ by extrapolating the growth curves of the infected cell cultures under selection $L(t)$, and without selection $T(t)$ to $t = 0$, defined as the time the samples in the infection plate were diluted into the detection plate (**Figure 3B**). This was done by fitting **Equation 9.1** to $L(t)$ and $T(t)$ for OD between ~0.02 and ~0.1. We note that in this case, the parameter $g$ reflects the growth rate in the detection plate, not the growth rates at which infection was performed.

In **Figures 3C** and **4B,** the calculated frequencies of lysogeny for each bacterial growth rate are plotted as a function of the MOI, $f_{\text{lysogeny}}(M)$.

### 10. Inferring the single-cell probability of lysogenization

#### 10.1. Model description

Following refs. [5,12,21,22], we assumed the following:

(1) Phage-cell encounters follow Poisson statistics. As a result, the single-cell MOI, $n$, is described by the following distribution.

$$P_n = \frac{(aM)^n e^{-aM}}{n!}, \tag{10.1}$$

where $M$ is the average MOI in the infection mixture, and $a$ is a scaling factor that accounts for the infection efficiency and the accuracy in measuring phage and cell concentrations.

(2) The probability of lysogenization, $Q_n$, is a function of the single-cell MOI, $n$. Unlike refs. [5,21,22] which assumed coinfection by at least $\mathrm{MOI}^*$ phages is required for lysogeny, we adopted the more general approach described in ref. [12], which allows for non-zero probability of lysogenization at $n < \mathrm{MOI}^*$:

$$Q_n = \begin{cases} 0 & \text{for } n = 0 \\ q_1 & \text{for } n = 1 \\ q_2 & \text{for } n = 2 \\ \ldots & \\ q_{\mathrm{MOI}^*} & \text{for } n \geq \mathrm{MOI}^*. \end{cases} \tag{10.2}$$

The observed frequency of lysogeny, $f_{\text{lysogeny}}$, as a function of the average MOI, $M$, is found by summing the product of $P_n$ and $Q_n$ over all possible values of $n$:

$$f_{\text{lysogeny}} = \sum_{n=0}^{\infty} P_n Q_n. \tag{10.3}$$

We scanned the value of $\mathrm{MOI}^*$ from 1 to 3, and found that $\mathrm{MOI}^* = 2$ best captured the lysogenization data by wild-type, replication-competent lambda phages in exponentially growing cells (consistent with refs. [5,22]). For $\mathrm{MOI}^* = 2$, **Equation 10.3** becomes:

$$f_{\text{lysogeny}} = P_1 q_1 + q_2 \sum_{n=2}^{\infty} P(n) = P_1 q_1 + (1 - P_0 - P_1) q_2. \tag{10.4}$$

**Equation 10.4** was used to fit the lysogeny data shown in **Figure 3C**.

#### 10.2. Incorporating the reduced frequency of lysogeny at high MOI

At some growth rates, we observed a reduction in the frequency of lysogeny at very high MOI (consistent with previous reports[21,22]). To capture this reduction, we introduced into **Equation 10.3** an additional repression term, $R_n$, which decreases the probability of lysogenization when a cell is infected at high $n$ values. We parameterized $R_n$ as an exponential decay with a rate of $k$:

$$R_n = \begin{cases} 1 & \text{for } 0 \leq n < \mathrm{MOI}^* \\ e^{-k(n - \mathrm{MOI}^*)} & \text{for } n \geq \mathrm{MOI}^*. \end{cases} \tag{10.5}$$

For up to $n = \mathrm{MOI}^*$, $R_n = 1$, thus the single-cell probability of lysogenization is simply $Q_n$. With this additional term, **Equation 10.3** becomes:

$$f_{\mathrm{lysogeny}} = \sum_{n=0}^{\infty} P_n Q_n R_n, \quad (10.6)$$

and **Equation 10.4** becomes:

$$f_{\mathrm{lysogeny}} = P_1 q_1 + q_2 \sum_{n=2}^{\infty} P_n R_n. \quad (10.7)$$

For $\mathrm{MOI}^* = 1$ (used to fit data in stationary cells in **Figure S13**), the frequency of lysogeny is:

$$f_{\mathrm{lysogeny}} = q_1 \sum_{n=1}^{\infty} P_n R_n. \quad (10.8)$$

##### 10.3. Model fitting

To capture the lysogeny-vs.-MOI curve measured at each bacterial growth rate, fitting was performed by minimizing the following objective function:

$$R(a, q_1, q_2, k) = \sum_{M} \left[\log f_{\mathrm{lysogeny}} - \log \hat{f}_{\mathrm{lysogeny}}\right]^2, \quad (10.9)$$

where $\hat{f}_{\mathrm{lysogeny}}$ are the model-predicted frequencies of lysogeny, and $f_{\mathrm{lysogeny}}$ are the experimentally measured values. Fitting was performed in logarithmic space because the MOI and the frequencies of lysogeny span several orders of magnitude. We also imposed a constraint of $q_1 \leq q_2$. Fitting results are shown in **Figures 3C** and **4B**, with the average MOI rescaled using the parameter $a$, as done in ref. [5].

We note that the best-fit values of the parameter $a$, when plotted as a function of the bacterial density, are in agreement with the theoretically predicted efficiency of phage adsorption[23] (**Figure S16A**). This agreement lends further credence to our measurements and the fitting procedure.

##### 10.4. Predicting the frequency of lysogeny as a function of both MOI and growth rates

To predict the parameters $a$, $q_1$, $q_2$, and $k$ at an arbitrary growth rate $g$, the best-fit values of these parameters at each sampled growth rate (obtained as described in **Section 10.3**) were used to fit the following phenomenological expressions:

$$\log a = \beta_2 g^2 + \beta_1 g + \beta_0,$$

$$\log q_1 = \beta_1 g + \beta_0,$$

$$\log q_2 = \begin{cases} \beta_1 g + \beta_0 & \text{for } g < g^* \\ \beta_2 g + (\beta_1 - \beta_2) g^* + \beta_0 & \text{for } g \geq g^*, \end{cases} \tag{10.10}$$

$$k = \begin{cases} 0 & \text{for } g < g_1 \\ -(g - g_1)(g - g_2) & \text{for } g_1 \leq g < g_2 \\ 0 & \text{for } g \geq g_2. \end{cases}$$

The results of these parameterizations are shown in **Figure S16B** (for $q_1$, also reproduced in **Figure 4C**), and the parameter values (with bootstrap standard errors) are shown in **Table S9**.

To predict the frequency of lysogeny as a function of both MOIs ($M$) and growth rates ($g$), we scanned the MOI between $10^{-3}$ and 100, and the growth rate between 0 and 1.25 $h^{-1}$. For each growth rate ($g$), the parameter values in **Table S9** and **Equations 10.10** were used to calculate the values of $a$, $q_1$, $q_2$, and $k$. These parameter values and **Equations 10.1**, **10.5**, and **10.7** were used to calculate the predicted frequency of lysogeny for different MOI values ($M$). The "fate diagram", depicting the frequency of lysogeny as a function of both MOI and growth rates, is shown in **Figure 4E**.

This model prediction was compared with the experimental data (**Figure 4E**). Here, the sampled data points (6 growth rates, each with 6 MOI values) were interpolated using the triangulation-based natural neighbor method and further smoothed using a 2D median filter, as done in ref. [24].

#### SUPPLEMENTARY FIGURES

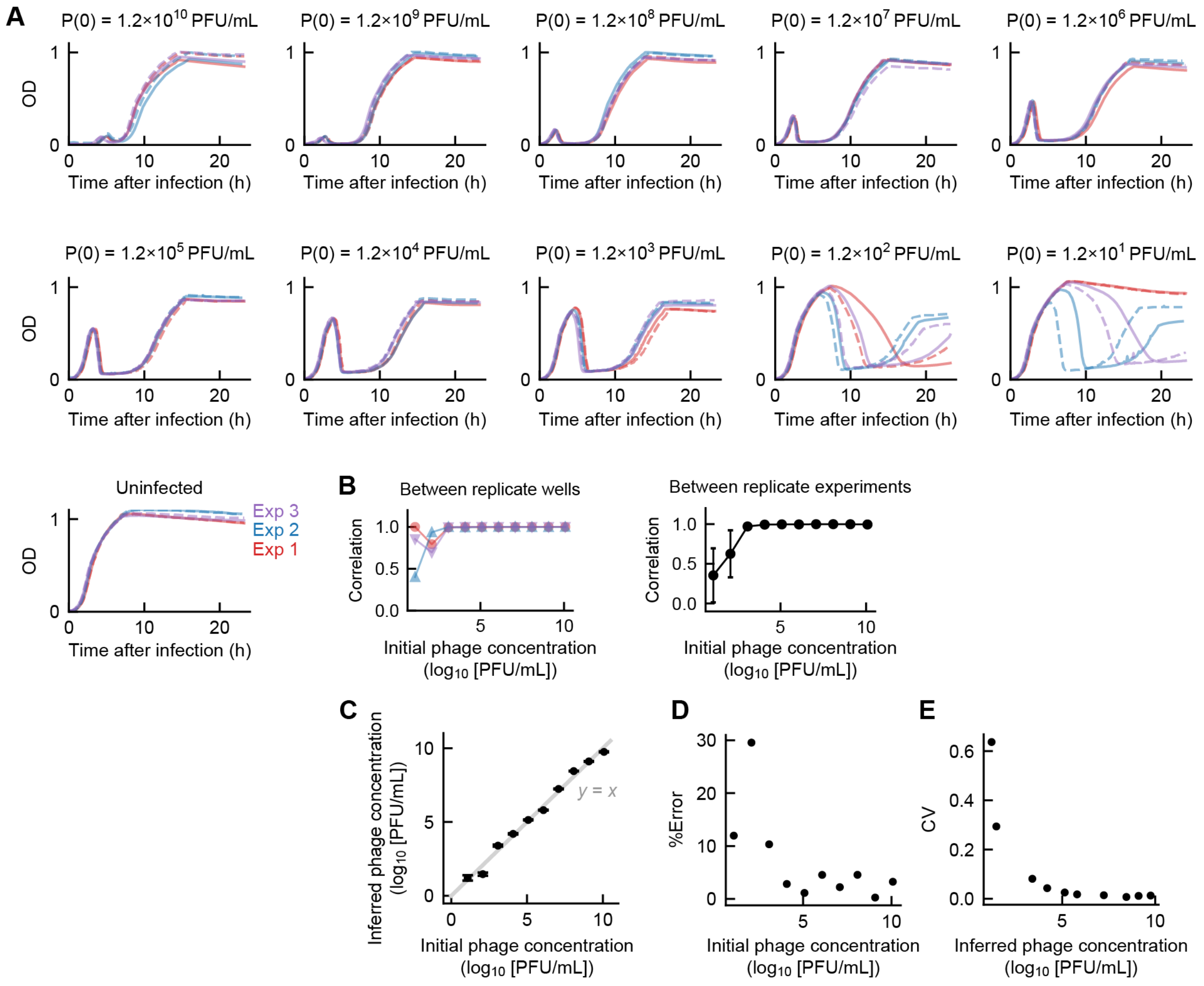

**Figure S1. Reproducibility of the bacterial growth dynamics and the OD-based phage counting method.**

**(A)** Representative growth curves of *E. coli* MG1655 cultures at 37ºC in LBM infected by $\lambda_{ts}$ at different concentrations (three experiments, different colors). For each initial phage concentration ($P(0)$) and each experiment, growth curves of two replicate wells (solid and dashed lines) are shown. The experimental protocol was described in ***Methods, Section 4***. **(B)** The reproducibility of growth curves within and between experiments. Left, Pearson correlation coefficients of the growth curves from two replicate cultures infected at a given initial phage concentration in each of the three experiments (as in Panel A). Right, similar to the left graph, calculated with bootstrapping ($N$ = 1000) from all replicate cultures across the three experiments. Error bar, standard error of the mean (SEM). **(C)** The error in inferring phage concentrations. For each initial phage concentration, we measured the lysis OD in six replicate cultures. For each lysis OD, we used three calibration curves (obtained in separate repeats of the experiment) to infer the corresponding initial phage concentration. Marker, average values. Error bars, SEM due to variations in the lysis OD and the calibration curves. Gray line, $y = x$. **(D)** The accuracy in inferring phage concentrations, calculated as $|\log_{10}$(Inferred concentration) - $\log_{10}$(Initial concentration)$|$ / $\log_{10}$(Initial concentration), using the values in Panel C. **(E)** The precision in inferring phage concentrations, calculated as the coefficients of variation (CV) of the values in Panel C.

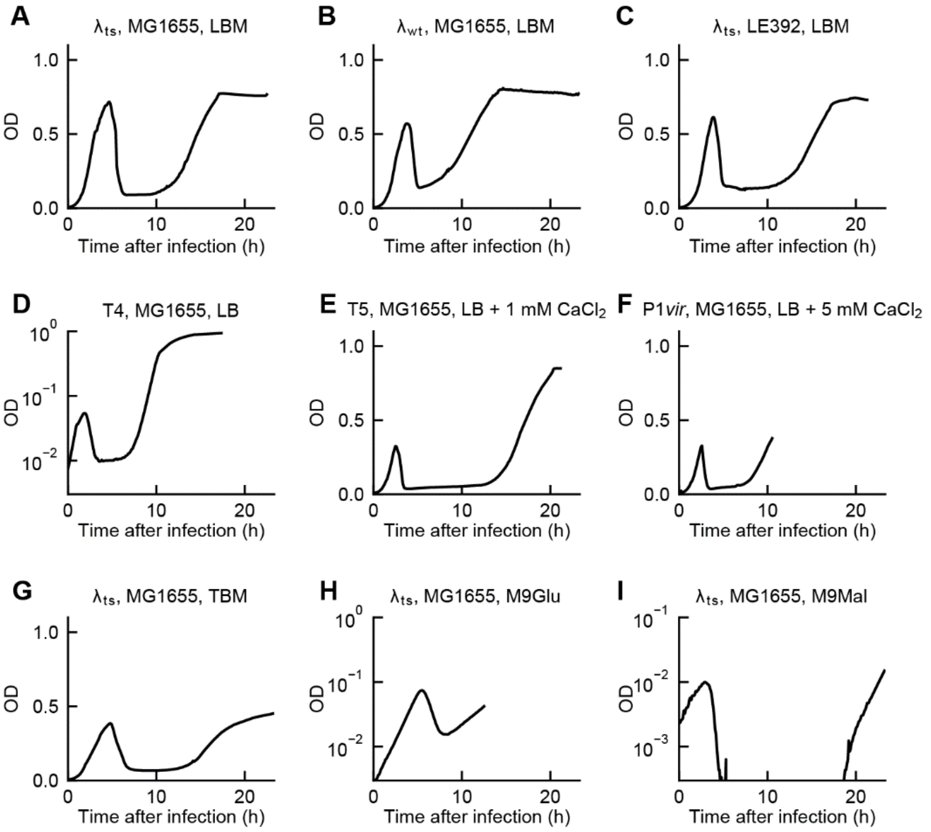

**Figure S2. Growth curves of infected bacterial cultures under different conditions.**

**(A-I)** Lines, growth curves of MG1655 cultures infected at 37°C. The initial phage concentration for each infection condition ranges between ≈ $1\times10^4$ PFU/mL and ≈ $7\times10^4$ PFU/mL. For infection by phage T4 (Panel D) and infections in minimal media (Panels H and I), the y-axes are displayed on a logarithmic scale.

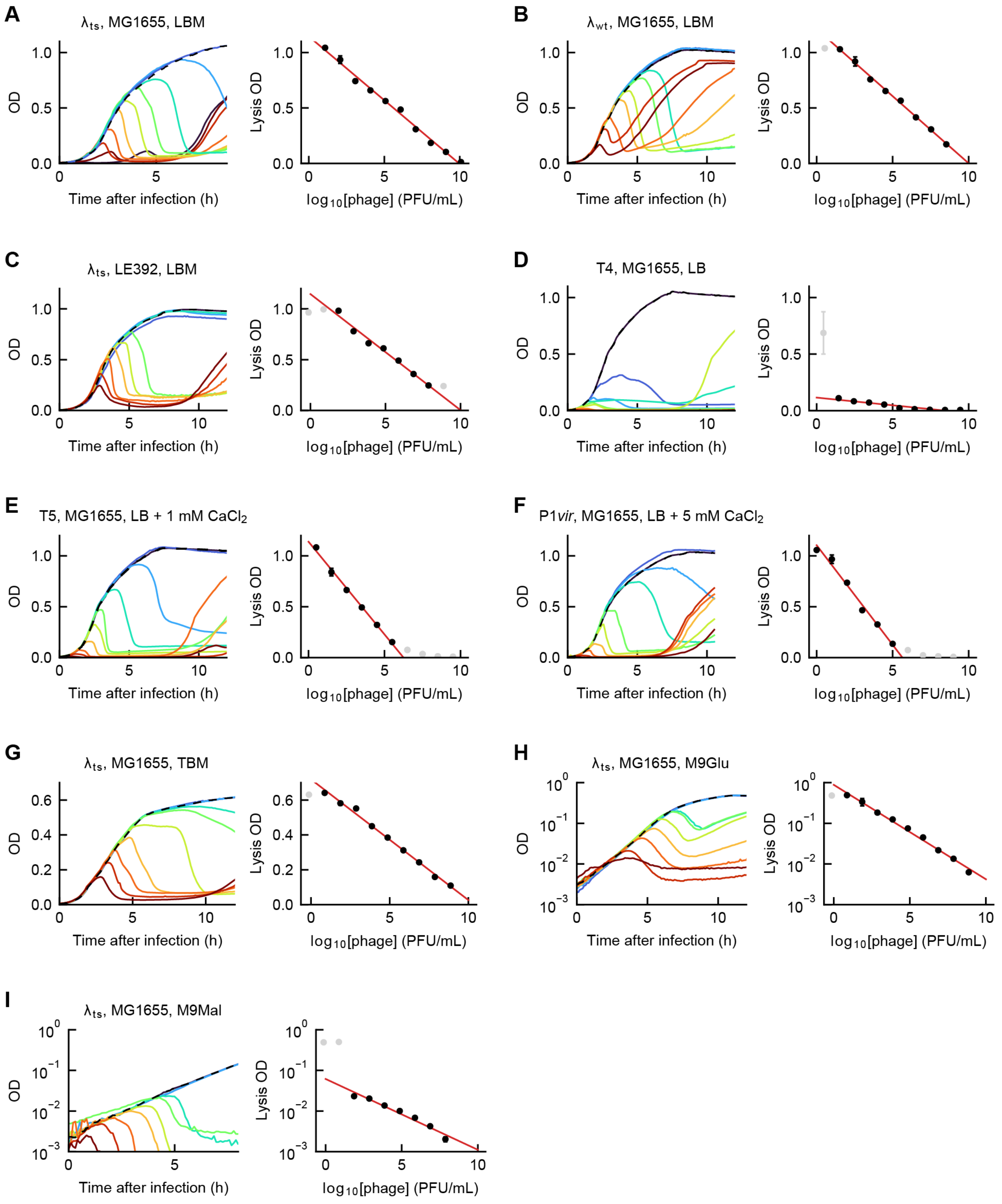

A
λts, MG1655, LBM
OD
Time after infection (h)
Lysis OD
log10[phage] (PFU/mL)
B
λwt, MG1655, LBM
C
λts, LE392, LBM
D
T4, MG1655, LB
E
T5, MG1655, LB + 1 mM CaCl2
F
P1vir, MG1655, LB + 5 mM CaCl2
G
λts, MG1655, TBM
H
λts, MG1655, M9Glu
I
λts, MG1655, M9Mal

**Figure S3. Calibration curves for counting phages under different infection conditions.**

The infection conditions in this figure correspond to those in **Figure S2**. In each panel, left: solid lines, growth curves of *E. coli* cultures infected at different phage concentrations (red to blue, high to low phage concentrations); dashed line, growth curve of an uninfected culture. Right: black markers, data used to fit the calibration curve. Error bars, SEM from culture replicates. Gray markers, data not used for fitting. Red line, linear fit ($y = k \cdot x + b$). For infections in minimal media (Panels H and I), the y-axes are displayed on a logarithmic scale. The fitting procedure was described in ***Methods, Sections 4.1*** and ***4.4***.

The range of phage concentrations and the fitted parameter values for each infection condition are as follows.

**(A)** From ≈ $1.2\times10^{1}$ PFU/mL to ≈ $1.2\times10^{10}$ PFU/mL, with $k = -0.115 \pm 0.002$, $b = 1.15 \pm 0.02$.

**(B)** From ≈ $3.4\times10^{0}$ PFU/mL to ≈ $3.4\times10^{8}$ PFU/mL, with $k = -0.121 \pm 0.004$, $b = 1.21 \pm 0.03$.

**(C)** From ≈ $7.4\times10^{-1}$ PFU/mL to ≈ $7.4\times10^{8}$ PFU/mL, with $k = -0.114 \pm 0.005$, $b = 1.15 \pm 0.03$.

**(D)** From ≈ $2.8\times10^{0}$ PFU/mL to ≈ $2.8\times10^{9}$ PFU/mL, with $k = -0.014 \pm 0.001$, $b = 0.12 \pm 0.01$.

**(E)** From ≈ $3.2\times10^{0}$ PFU/mL to ≈ $3.2\times10^{9}$ PFU/mL, with $k = -0.19 \pm 0.01$, $b = 1.13 \pm 0.03$.

**(F)** From ≈ $1.2\times10^{0}$ PFU/mL to ≈ $1.2\times10^{9}$ PFU/mL, with $k = -0.20 \pm 0.01$, $b = 1.10 \pm 0.04$.

**(G)** From ≈ $7.4\times10^{-1}$ PFU/mL to ≈ $7.4\times10^{8}$ PFU/mL, with $k = -0.069 \pm 0.002$, $b = 0.72 \pm 0.01$.

**(H)** From ≈ $7.4\times10^{-1}$ PFU/mL to ≈ $7.4\times10^{8}$ PFU/mL, with $k = -0.23 \pm 0.01$, $b = -0.05 \pm 0.06$.

**(I)** From ≈ $7.4\times10^{-1}$ PFU/mL to ≈ $7.4\times10^{8}$ PFU/mL, with $k = -0.17 \pm 0.01$, $b = -1.21 \pm 0.07$.

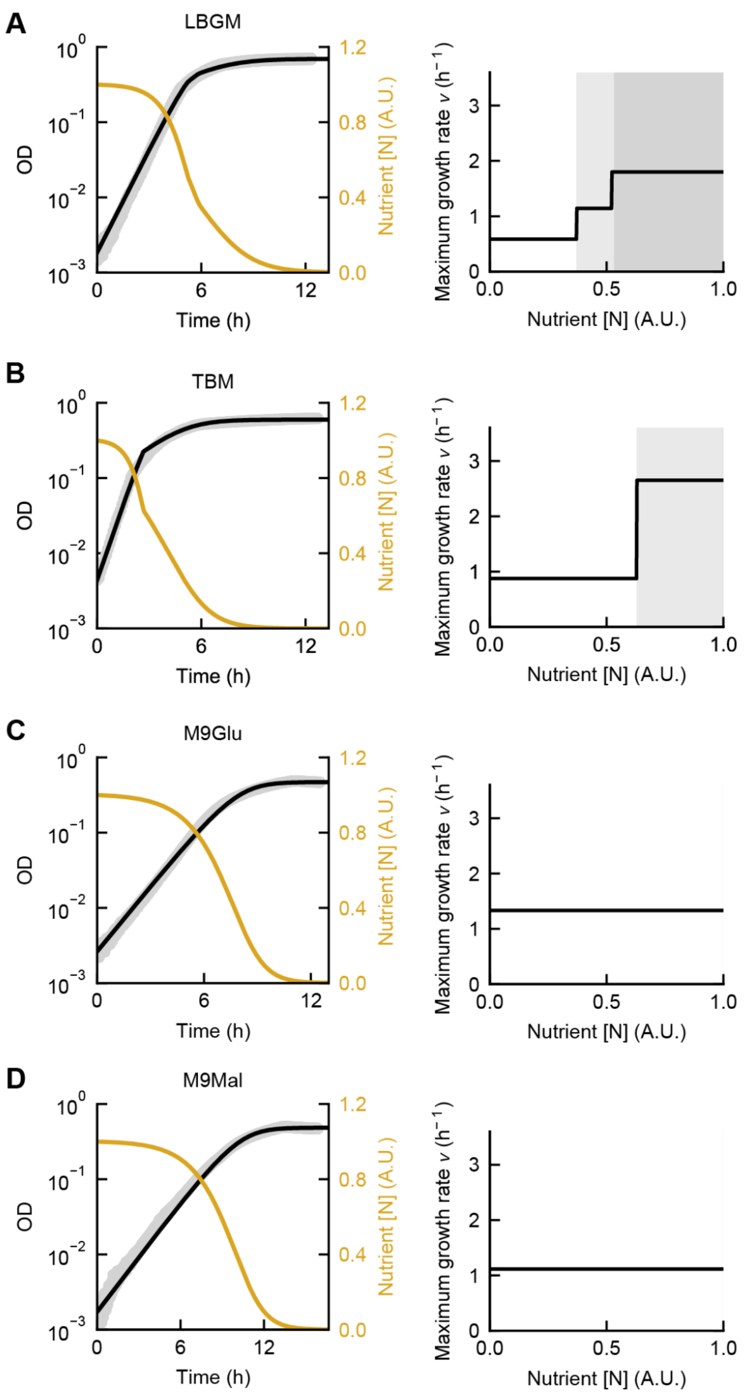

**Figure S4. Parameterization of bacterial growth in (A) LBGM, (B) TBM, (C) M9Glu, and (D) M9Mal.**

MG1655 cells were grown in different growth media at 37°C. In each panel, left, a model describing nutrient-dependent growth (black) captures the measured OD curves of uninfected cultures (gray); gold, the inferred time-dependent nutrient abundance. Right, the maximum growth rate $v$ (black) at different stages of nutrient consumption (white and gray shading). The fitted parameters are shown in **Tables S2, S3** and **S7**. The modeling for cell growth was described in ***Methods, Section 6.2***.

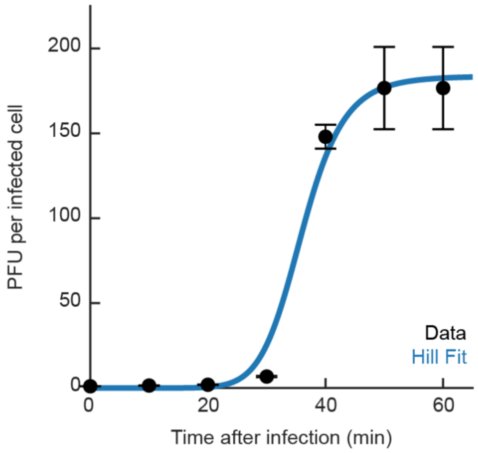

**Figure S5. One-step growth curve for infection of MG1655 by $\lambda_{ts}$.**

Infection was performed as described in ref. [5]. Briefly, cells were cultured in LBMM at 37°C, infected at MOI ≈ 0.1, diluted into pre-warmed LBGM, and grown at 37°C with aeration. At each sampled time point, infected cells were aliquoted and immediately assayed for the phage concentration using the plaque formation assay. The numbers of PFU per infected cell were calculated by normalizing the measured phage concentrations by the average at 0 and 10 minutes. Markers, data; error bars, SEM between plating replicates. Blue curve, Hill fit ($y = (ax^n)/(k^n + x^n)$). Burst size (estimated using the parameter $a$), $184 \pm 28$ PFU per cell. Latent period (estimated using the parameter $k$), $36.0 \pm 3.3$ min. For comparison with model-inferred values, see ***Tables S5*** and ***S6**.*

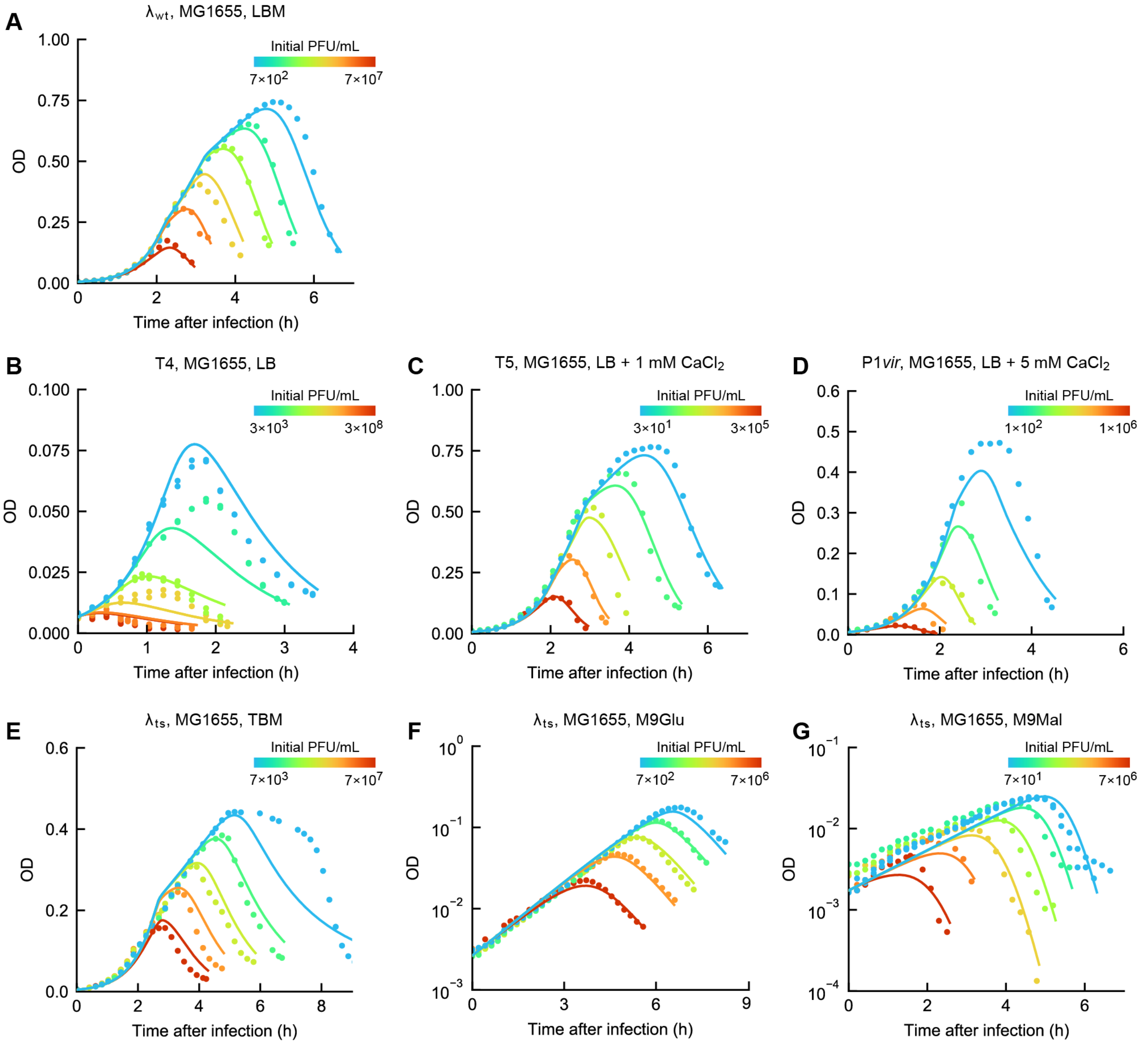

**Figure S6. Model fitting for the bacterial growth dynamics under different infection conditions.**

Colored markers, data from infection at different initial phage concentrations. Colored lines, model fits. The fitting procedure was described in ***Methods, Sections 6.1–6.5***. The fitted parameters are shown in **Table S6**.

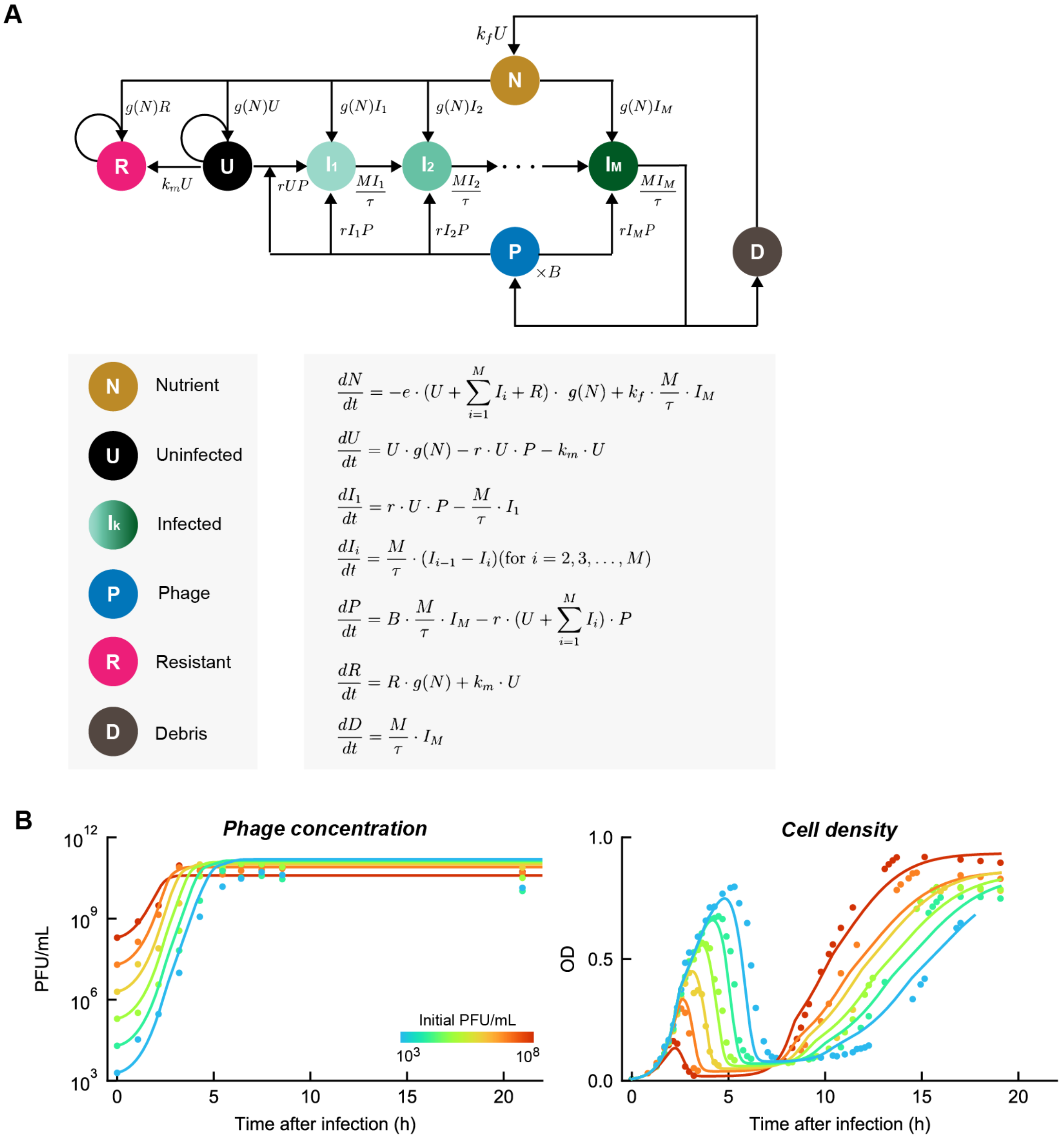

**Figure S7. Modeling bacterial recovery following phage infection.**

**(A)** A model with a transition from phage-sensitive to resistant cells captures the observed recovery in bacterial growth. Circles, species tracked by the model. Arrows, transitions between species. The transition rates are indicated next to the corresponding arrows. **(B)** Model fitting for the measured phage concentrations (left) and cell densities (right) over time. Colored markers, data from infection at different initial phage concentrations (reproduced from **Figure 2D**). Colored lines, model fits. The fitting procedure was described in ***Methods, Section 6.6***. The fitted parameters are as follows. $k_m = (6.6 \pm 0.8) \times 10^{-7}$ min$^{-1}$, $k_f = 0.797 \pm 0.005$. Parameters related to cell growth are shown in **Table S4**. The remaining parameters ($r$ and $B$) are shown in **Table S6**.

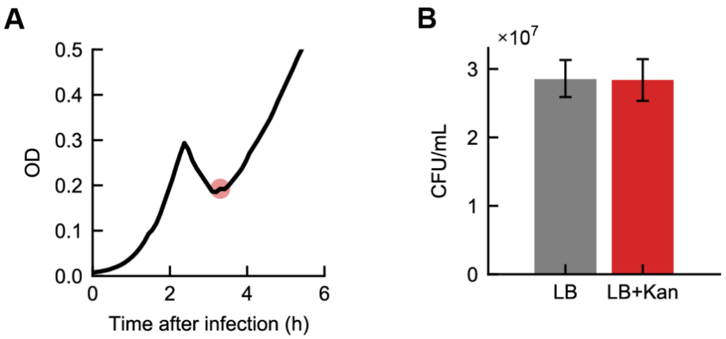

**Figure S8. Quantifying the proportion of lysogens among surviving cells in phage-infected cultures.**

MG1655 cells were infected by temperate phage $\lambda_{wt}$, as described in ***Methods, Section 7***. **(A)** The growth curve following infection at an initial phage concentration of ≈ $2 \times 10^7$ PFU/mL. Red circle, the point at which the culture was extracted for colony plating. **(B)** Gray bar, total cell concentration ($(2.85 \pm 0.26) \times 10^7$ CFU/mL), obtained by plating on non-selective LB agar plates. Red bar, concentration of lysogenic cells ($(2.84 \pm 0.31) \times 10^7$ CFU/mL), obtained by plating on LB agar plates supplemented with 50 μg/mL kanamycin. Error bars, SEM from dilution and plating replicates.

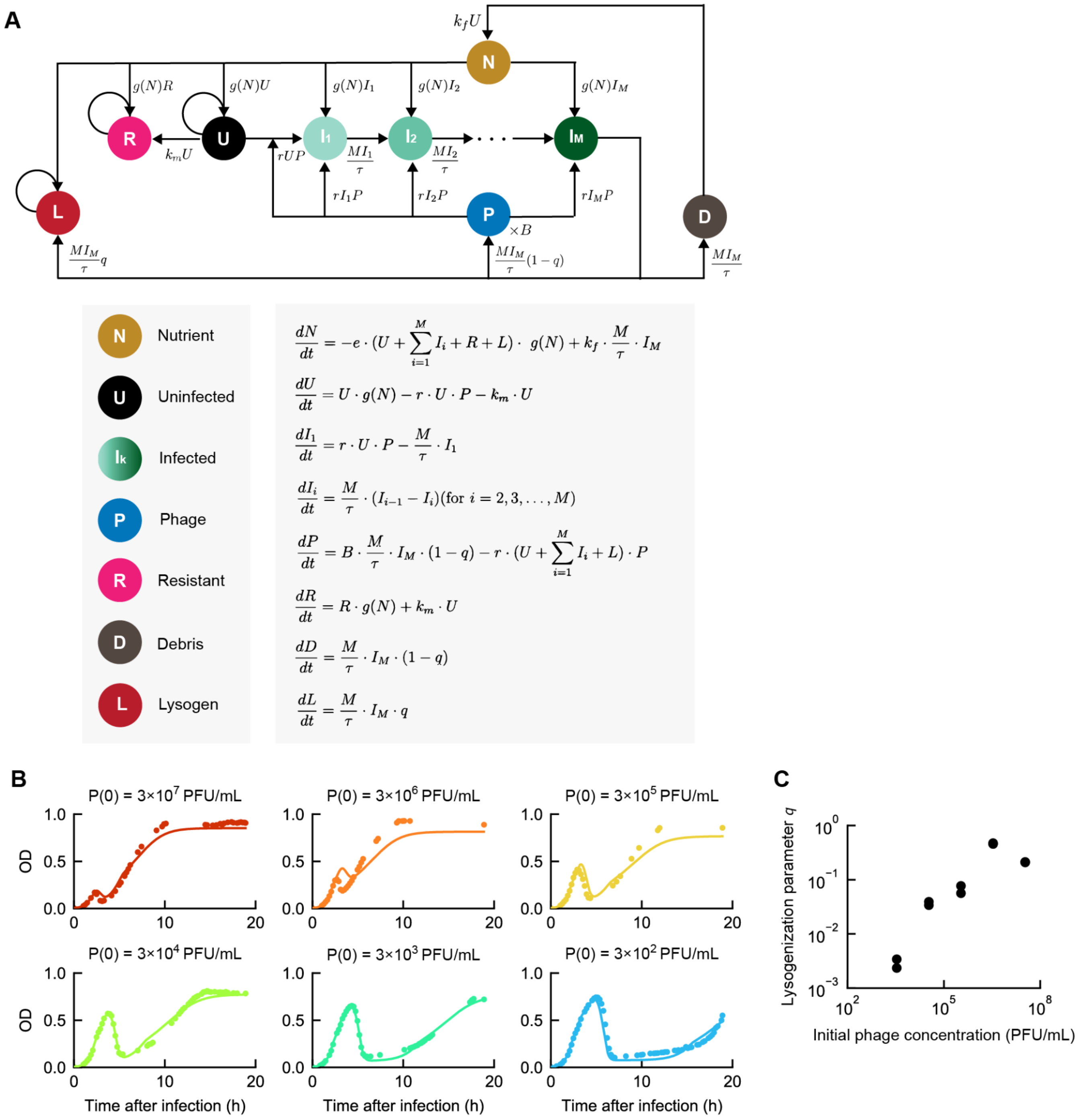

**Figure S9. Modeling infection by temperate phages.**

**(A)** Model schematics and equations. Circles, species tracked by the model. Arrows, transitions between species. The transition rates are indicated next to the corresponding arrows. **(B)** Model fitting for the measured cell densities over time. Colored markers, data from infection at different initial phage concentrations. Colored lines, model fits. The fitting procedure was described in ***Methods, Section 6.7***. **(C)** The lysogenization parameter $q$, inferred from the fits in Panel B, as a function of the initial phage concentration.

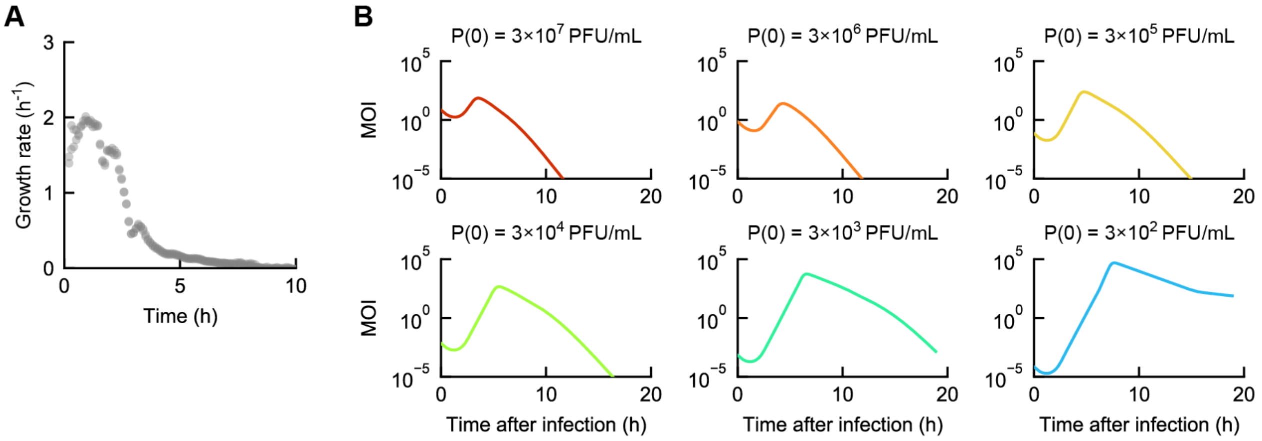

**Figure S10. Changes in growth rate and MOI of an infected bacterial culture over time.**

**(A)** The instantaneous growth rate of an uninfected MG1655 culture in LBM medium at 37°C (prepared as described in ***Methods, Section 4***). The growth rate $g$ was calculated by fitting **Equation 9.1** to the OD curve over a time window of 20 minutes. **(B)** The average MOI of the cell cultures, each infected by phage $\lambda_{wt}$ at different initial concentrations ($P(0)$). The time-dependent MOI was calculated using the model-inferred phage concentrations and cell densities (as depicted in **Figure S9B**). The model was described in ***Methods, Section 6.7***.

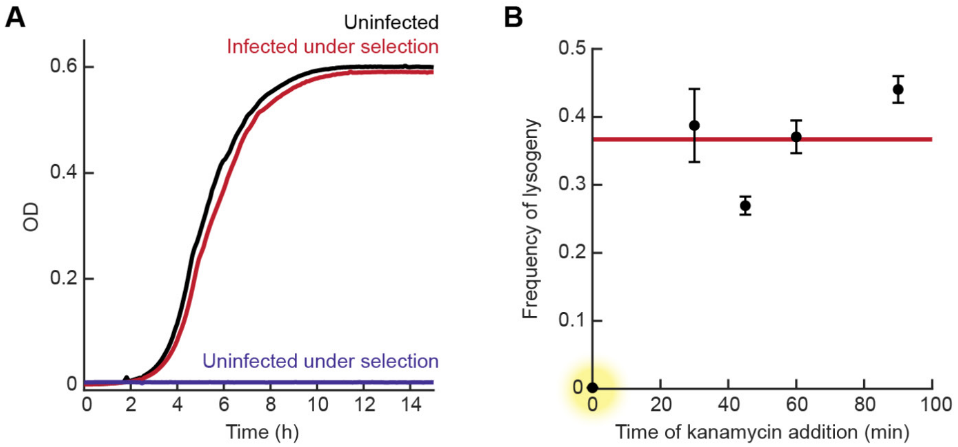

**Figure S11. Using kanamycin to select for lysogenic cells.**

**(A)** The growth curves of MG1655 without kanamycin selection (black) and under selection of 50 μg/mL kanamycin (violet), and of lysogenic cells (MG1655 $\lambda_{ts}$) under selection (red). All cultures were inoculated using similar concentrations of cells, and grown in LBGM at 30°C. **(B)** The frequency of lysogeny (measured as described in ***Methods, Section 9.3***) as a function of the time when kanamycin was added after infection. MG1655 cells were infected by $\lambda_{ts}$ at MOI ≈ 5 (using the same protocol described in **Figure S5**), and incubated at 30°C. At different times, 50 μg/mL kanamycin was added to the culture. Markers, data; error bars, SEM from technical replicates. Red line, average of the values at 30, 45, 60, and 90 minutes. When kanamycin was added immediately after infection (yellow highlight), the measured frequency was only 0.002.

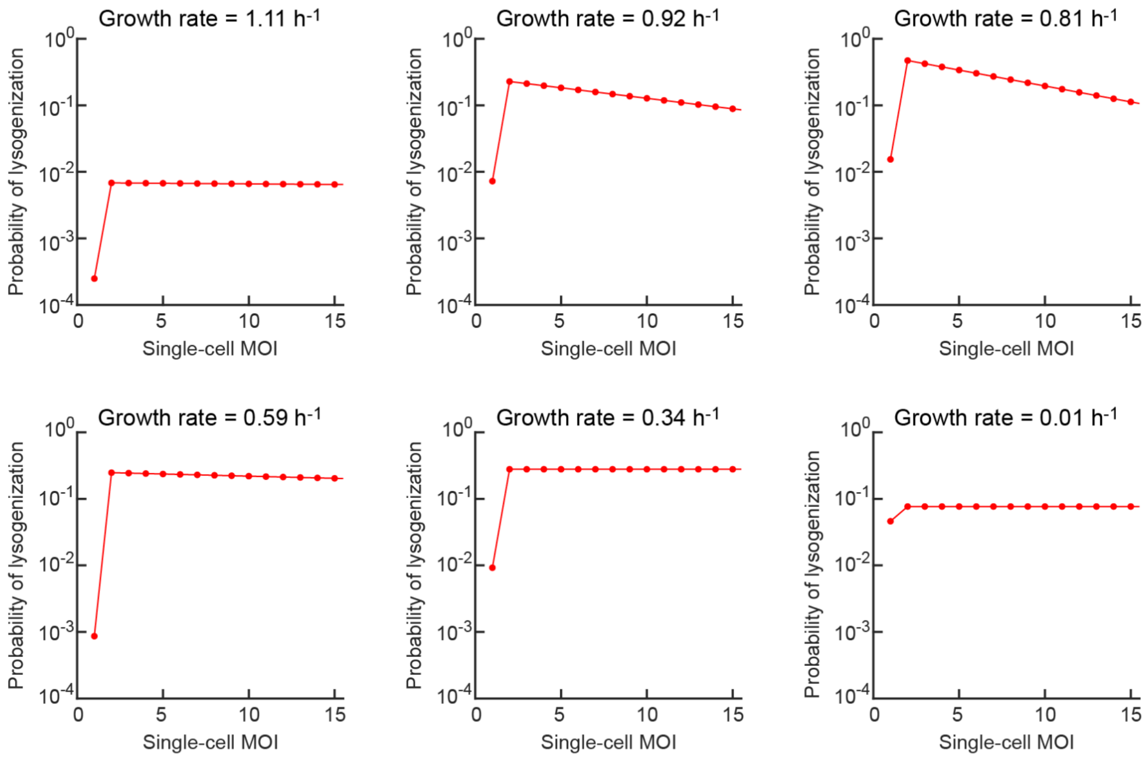

**Figure S12. The inferred single-cell MOI response curve at different growth rates.**

MG1655 cells at different growth rates were infected by $\lambda_{ts}$ as described in ***Methods, Section 9.*** Markers, the probability of lysogenization (inferred using the product $Q_n R_n$, as described in ***Methods, Section 10***) as a function of the single-cell MOI ($n$).

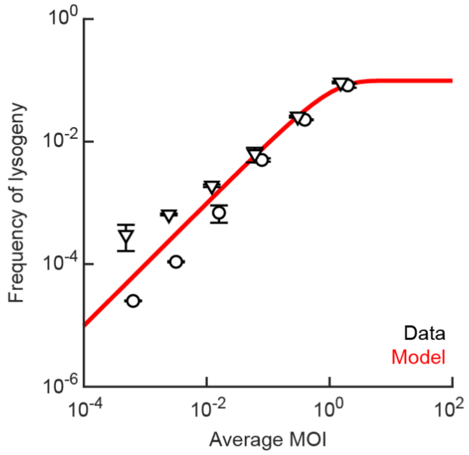

**Figure S13. Fitting the frequency of lysogeny in stationary cells, using MOI* = 1.**

MG1655 cells in stationary phase were infected by $\lambda_{ts}$ as described in ***Methods, Section 9*.** Circles and triangles, data obtained in two independent runs of the experiment (reproduced from **Figure 4B**); error bars, SEM. Red line, model fit using **Equation 10.8**, which assumes MOI* = 1 (in contrast to the model shown in **Figure 4B**, fitted using **Equation 10.7**, which assumes MOI* = 2).

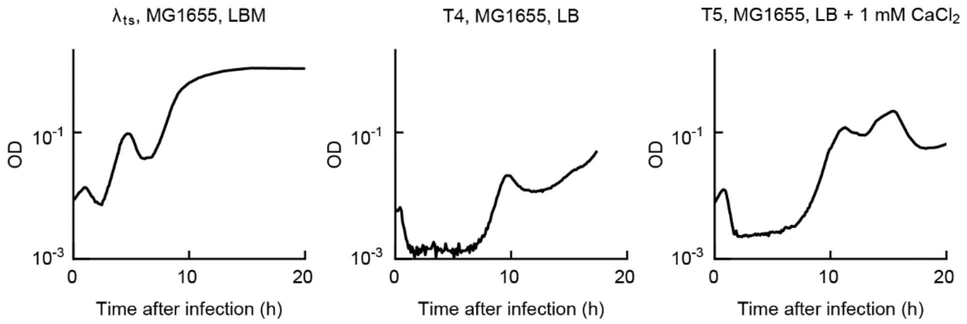

**Figure S14. Multiple cycles of growth and lysis following phage infection.**

Lines, growth curves of MG1655 cells when infected by phages $\lambda_{ts}$, T4, and T5 at MOI ≈ 1000. Infection mixtures were incubated at 37°C in LB medium (supplemented with 10 mM $MgSO_4$ for $\lambda_{ts}$, or 1 mM $CaCl_2$ for T5). The experimental protocol was described in ***Methods, Section 4***.

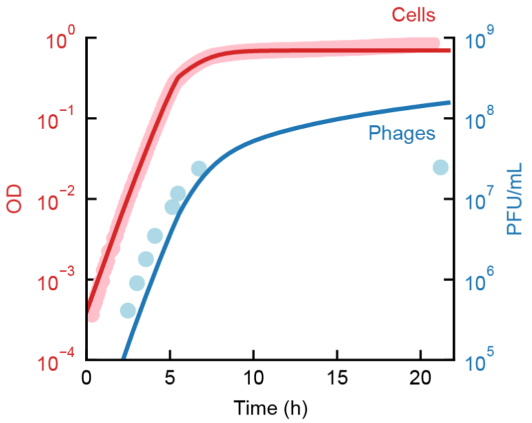

**Figure S15. Modeling spontaneous induction without the dependence of induction rate on the bacterial growth rate fails to capture the data.**

The densities of bacteria (red) and free phages (blue) during growth of lysogens. MG1655 $\lambda_{ts,}$ was grown at 30$^o$C in LBM supplemented with 0.2% glucose. Markers, data (reproduced from **Figure 4F**). Lines, model fits. Here, the model assumes the induction rate is independent of the bacterial growth rate (in contrast to the model shown in **Figure 4F**). The fitting procedure was described in ***Methods, Section 8.2***. The fitted spontaneous induction rate $k_i$ is $(1.9 \pm 0.0) \times 10^{-4}$ per hour.

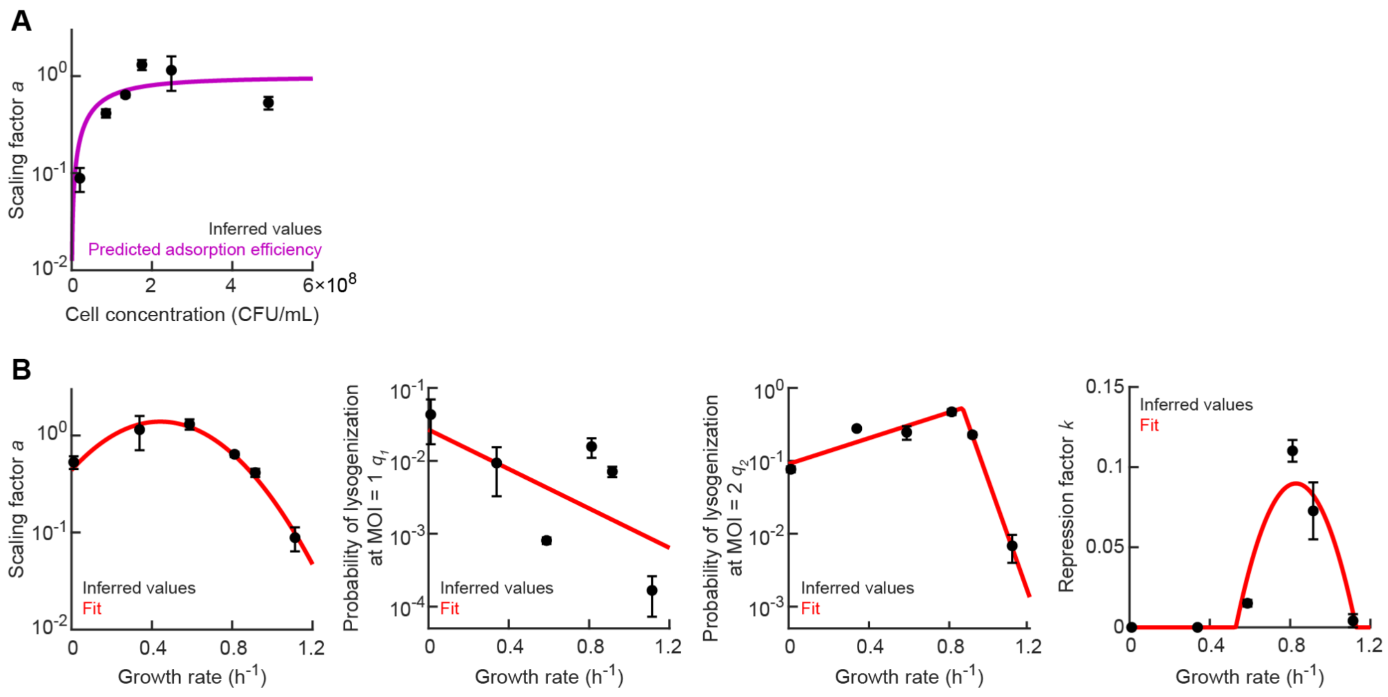

**Figure S16. Parameterization of the frequency of lysogeny as a function of growth rate.**

**(A)** The MOI-scaling factor $a$ is consistent with the efficiency of phage adsorption. Markers, values of the parameter $a$, inferred as described in ***Methods, Section 10***, following infection at different bacterial densities (***Methods, Section 9***); error bars, SEM from two independent runs of the experiment. Purple curve, prediction by the model in ref. [23] for maltose-containing medium at 30°C. **(B)** The parameters $a$, $q_1$, $q_2$, and $k$, inferred as described in ***Methods, Section 10***, as a function of the bacterial growth rate upon infection. Markers, inferred values; error bars, SEM from two independent runs of the experiment (Values for $a$ are reproduced from Panel A). Red lines, parameterization using **Equation 10.10** (see **Table S9** for the parameter values).

### SUPPLEMENTARY TABLES

**Table S1: Bacterial and phage strains used in this study**

| Strain | Description | Source |
|---|---|---|
| **BACTERIA** | | |
| MG1655 | Wild-type *E. coli* | Lab stock |
| LE392 | *glnV* (*supE*44), *tryT* (*supF*58); amber suppressor[25]. | Lab stock |
| **PHAGE** | | |
| $\lambda_{ts}$ | λ *cI*857 *bor*::*kan*$^R$<br>Temperature-sensitive; obligately lytic at 37°C and above[1]. | Lab stock |
| $\lambda_{wt}$ | λ $cI_{wt}$ *bor*::*kan*$^R$<br>Wild-type. | Lab stock |
| T4 | | Coli Genetic Stock Center |
| T5 | | Coli Genetic Stock Center |
| P1*vir* | Virulent mutant | Lab stock |

**Table S2: Fitted growth parameters in single-carbon growth media ($X = 1$)**

| Parameter | Description | Mean $\pm$ SEM | |
|---|---|---|---|
| | | **M9Glu** | **M9Mal** |
| $v_1$ | Maximum growth rate | $0.022 \pm 0.000$ min$^{-1}$ | $0.019 \pm 0.000$ min$^{-1}$ |
| $K_1$ | Affinity constant | $1.0 \pm 0.0$ | $1.0 \pm 0.0$ |
| $e$ | Conversion efficacy parameter | $2.1 \pm 0.0$ | $2.1 \pm 0.0$ |

**Table S3: Fitted growth parameters in TBM ($X = 2$)**

| Parameter | Description | Mean $\pm$ SEM |
|---|---|---|
| $v_1$ | Maximum growth rate in phase 1 | $0.044 \pm 0.003$ min$^{-1}$ |
| $v_2$ | Maximum growth rate in phase 2 | $0.015 \pm 0.001$ min$^{-1}$ |
| $K_1$ | Affinity constant for substrate in phase 1 | $0.72 \pm 0.08$ |
| $K_2$ | Affinity constant for substrate in phase 2 | $0.88 \pm 0.05$ |
| $N_1$ | The nutrient concentration at which phase 1 ends | $0.63 \pm 0.04$ |
| $e$ | Conversion efficacy parameter | $1.68 \pm 0.01$ |

**Table S4: Fitted growth parameters in LBM ($X = 3$)**

| Parameter | Description | Mean $\pm$ SEM |
|---|---|---|
| $v_1$ | Maximum growth rate in phase 1 | $0.054 \pm 0.002$ min$^{-1}$ |
| $v_2$ | Maximum growth rate in phase 2 | $0.018 \pm 0.002$ min$^{-1}$ |
| $v_3$ | Maximum growth rate in phase 3 | $0.008 \pm 0.001$ min$^{-1}$ |
| $K_1$ | Affinity constant for substrate in phase 1 | $0.92 \pm 0.07$ |
| $K_2$ | Affinity constant for substrate in phase 2 | $0.67 \pm 0.08$ |
| $K_3$ | Affinity constant for substrate in phase 3 | $0.47 \pm 0.09$ |
| $N_1$ | The nutrient concentration at which phase 1 ends | $0.57 \pm 0.06$ |
| $N_2$ | The nutrient concentration at which phase 2 ends | $0.52 \pm 0.04$ |
| $e$ | Conversion efficacy parameter | $0.94 \pm 0.00$ |

**Table S5: Fitted parameters for the latent period**

| Infecting phage | Growth media | $M$ | $b$ (Mean $\pm$ SEM) | $k$ (Mean $\pm$ SEM) |
|---|---|---|---|---|
| $\lambda_{ts}$ | LBM | 5 | $21.0 \pm 1.9$ min | $0.15 \pm 0.01$ |
| $\lambda_{ts}$ | TBM | 2 | $28.7 \pm 3.6$ min | $0.22 \pm 0.03$ |
| $\lambda_{ts}$ | M9Glu | 2 | $84.5 \pm 0.0$ min | $0$ |
| $\lambda_{ts}$ | M9Mal | 4 | $44.1 \pm 0.0$ min | $0$ |
| $\lambda_{wt}$ | LBM | 5 | $32.2 \pm 2.7$ min | $0.10 \pm 0.01$ |
| T4 | LB | 2 | $44.4 \pm 0.0$ min | $0$ |
| T5 | LB + 1 mM $CaCl_2$ | 4 | $31.9 \pm 4.2$ min | $0.09 \pm 0.09$ |
| P1 | LB + 5 mM $CaCl_2$ | 5 | $20.4 \pm 8.2$ min | $0.32 \pm 0.15$ |

**Table S6: Fitted values for the encounter rate and the burst size**

| Infecting phage | Growth media | $r$ (Mean $\pm$ SEM) | $b$ (Mean $\pm$ SEM) |
|---|---|---|---|
| $\lambda_{ts}$ | LBM | $(9.9 \pm 0.1) \times 10^{-12}$ mL·min$^{-1}$ | $207 \pm 4$ |
| $\lambda_{ts}$ | TBM | $(6.7 \pm 0.3) \times 10^{-12}$ mL·min$^{-1}$ | $370 \pm 22$ |
| $\lambda_{ts}$ | M9Glu | $(9.4 \pm 0.1) \times 10^{-11}$ mL·min$^{-1}$ | $114 \pm 2$ |
| $\lambda_{ts}$ | M9Mal | $(6.65 \pm 0.01) \times 10^{-10}$ mL·min$^{-1}$ | $83 \pm 0$ |
| $\lambda_{wt}$ | LBM | $(5.1 \pm 0.1) \times 10^{-11}$ mL·min$^{-1}$ | $72 \pm 2$ |
| T4 | LB | $(4.57 \pm 0.01) \times 10^{-8}$ mL·min$^{-1}$ | $15 \pm 0$ |
| T5 | LB + 1 mM $CaCl_2$ | $(9.58 \pm 0.03) \times 10^{-10}$ mL·min$^{-1}$ | $18 \pm 0$ |
| P1 | LB + 5 mM $CaCl_2$ | $(6.7 \pm 0.0) \times 10^{-10}$ mL·min$^{-1}$ | $35 \pm 0$ |

**Table S7: Fitted growth parameters in LBGM**

| Parameter | Description | Mean $\pm$ SEM |
|---|---|---|
| $v_1$ | Maximum growth rate in phase 1 | $0.030 \pm 0.001$ min$^{-1}$ |
| $v_2$ | Maximum growth rate in phase 2 | $0.019 \pm 0.002$ min$^{-1}$ |
| $v_3$ | Maximum growth rate in phase 3 | $0.010 \pm 0.001$ min$^{-1}$ |
| $K_1$ | Affinity constant for substrate in phase 1 | $0.71 \pm 0.07$ |
| $K_2$ | Affinity constant for substrate in phase 2 | $0.77 \pm 0.09$ |
| $K_3$ | Affinity constant for substrate in phase 3 | $0.73 \pm 0.06$ |
| $N_1$ | The nutrient concentration at which phase 1 ends | $0.52 \pm 0.04$ |
| $N_2$ | The nutrient concentration at which phase 2 ends | $0.37 \pm 0.09$ |
| $e$ | Conversion efficacy parameter | $1.439 \pm 0.003$ |

**Table S8: Parameterization for the spontaneous induction rate as a function of growth rate**

| Parameter | Mean $\pm$ SEM |
|---|---|
| $k$ | $(1.9 \pm 0.0) \times 10^{-4}$ |
| $b$ | $0.0 \pm 0.0$ hr$^{-1}$ |

**Table S9: Parameterization for the frequency of lysogeny as a function of MOI and growth rate**

| Parameter | Mean $\pm$ SEM |
|---|---|
| $\boldsymbol{a}$ | |
| $\beta_2$ | $-5.89 \pm 0.87$ |
| $\beta_1$ | $5.21 \pm 1.02$ |
| $\beta_0$ | $-0.82 \pm 0.31$ |
| $\boldsymbol{q_1}$ | |
| $\beta_1$ | $-3.11 \pm 1.33$ |
| $\beta_0$ | $-3.62 \pm 0.85$ |
| $\boldsymbol{q_2}$ | |
| $\beta_1$ | $2.04 \pm 0.00$ |
| $\beta_0$ | $-2.04 \pm 0.00$ |
| $\beta_2$ | $17.89 \pm 0.00$ |
| $g^*$ | $0.87 \pm 0.00$ |
| $\boldsymbol{k}$ | |
| $g_1$ | $0.53 \pm 0.06$ |
| $g_2$ | $1.13 \pm 0.06$ |
